# Isogenic reciprocal grafts with transgenic HaHB11 plants dissect shoot and root contributions to yield in field-grown soybean: a multi-omic study

**DOI:** 10.64898/2026.08.15.744928

**Authors:** Jesica Raineri, Gianni Rositto, Agustín Lucas Arce, María Elena Otegui, Raquel Lía Chan

## Abstract

Soybean must coordinate root and shoot signals to optimize yield. Grafting is a powerful tool to study this communication. However, most studies compare contrasting genotypes and cannot separate genotype from graft combination effects. Here we used isogenic soybean lines to dissect root and shoot contributions in the field. These lines differ from controls in a single gene, either HaHB11 or HaHB4, two sunflower HD-Zip I transcription factors associated with increased grain number. Unexpectedly, heterografted plants outperformed homografts in several yield-related traits, an effect not previously documented in soybean. This advantage was reproduced with both HaHB11 and HaHB4 scions, suggesting the effect is not gene-specific. Under non-stress conditions the scion governed yield-related traits, particularly pod number, as well as the leaf transcriptome, whereas both organs left subtle metabolic signatures. The root contribution was minor and confined to the R6–R7 transition, where it was specific to HaHB11. The highest-yielding combination was a control rootstock with an HaHB11 scion (CH11), which increased grain number by ∼30% over the best homograft. CH11 showed higher stomatal conductance and lower leaf temperature; yet CH11 and HaHB11 homografts were remarkably similar, sharing higher stomatal density, differing in only four leaf-expressed genes, and lacking a metabolomic signature. Thus, under non-stress conditions, soybean grain number is governed by the scion and the graft combination, and accompanied by early physiological differences rather than by leaf molecular reprogramming.

## Introduction

Plant growth and yield arise from the integration of shoot photosynthesis and root-mediated acquisition of water and nutrients. These processes are tightly interconnected through source-sink relationships and long-distance signaling pathways that coordinate whole-plant responses (Passioura, 1983; Tollenaar and Aguilera, 1992).

Soybean (*Glycine max (L.) Merr*.) is one of the most important crops worldwide, providing a major source of plant protein and vegetable oil (Lam et al., 2010). Despite this, unlike the cereal crops at the center of the Green Revolution, soybean has historically received comparatively less breeding attention and investment than maize or wheat, partly because its autogamous reproductive biology offers weaker incentives for private research investment (Fang et al., 2026; Fuglie et al., 2016). Even so, breeding programs have progressively selected varieties with improved seed number, photosynthetic capacity and harvest index (Jin et al., 2010; Rogers et al., 2015; Koester et al., 2014, 2016). This yield improvement has largely relied on the selection of aboveground traits, while the contribution of root systems to yield formation has been comparatively overlooked. This was partly due to the difficulty of phenotyping roots *in situ*, resulting in modern cultivars whose belowground traits probably have been indirectly co-selected rather than optimized (Cao et al., 2019; Koester et al., 2016; Li et al., 2017b).

Grafting provides a powerful experimental approach to unravel root and shoot contributions by combining genetically distinct scions and rootstocks in a single plant. Historically, grafting has been used for many plant species to propagate and to improve agronomic performance, including yield, stress tolerance and disease resistance (Mudge et al., 2009; Lee et al., 2010). More recently, grafting has emerged as a tool to study long-distance signaling, enabling the analysis of mobile molecules such as hormones, proteins, and RNAs that coordinate development and stress responses across organs (Augstein and Melnyk, 2025; Loupit et al., 2023).

In soybeans, grafting has been widely used to investigate root-shoot interactions underlying yield. A series of studies using cultivars released over different decades demonstrated that high-yielding rootstocks can enhance photosynthetic capacity and yield components of scions, suggesting that root function may limit further yield improvement (Li et al., 2017a; Li et al., 2017b; Du et al., 2019). Conversely, reciprocal experiments showed that elite scions can reprogram root traits, leading to reduced root biomass and altered root-shoot allocation, revealing an uncoordinated improvement between aboveground and belowground traits during breeding (Li et al., 2019). Grafting studies under abiotic stress further support the role for roots in determining plant performance under stress conditions (Li et al., 2021). In addition, molecular analyses have provided evidence of long-distance signaling between soybean roots and shoots. It was reported that root genotype can control leaf gene expression within hours (Kohlhase et al., 2025), and root-controlled nitrogen status can influence shoot photosynthesis and metabolism (He et al., 2021).

Despite these advances, previous grafting studies in soybean have primarily relied on genetically distinct cultivars or near-isogenic lines that differ in large genomic regions. This approach limits the ability to attribute observed phenotypic and molecular differences to specific genes. Moreover, most studies have focused either on agronomic traits or on molecular responses, but rarely integrate both levels under field conditions. Consequently, the contribution of individual genes to root–shoot communication and yield formation remains largely unresolved.

In addition, grafting itself can influence plant performance: combining tissues of different plants (even with the same genotype, homografts) may alter characteristics relative to self-grafted (a plant grafted to itself) controls (Cookson et al., 2014). Distinguishing this combination effect from the contribution of each organ’s genotype therefore requires reciprocal designs that include both homo- and heterografts.

In several species, including tomato, melon, Arabidopsis, and some woody plants, the differences between homo- and heterografted plants have been studied mainly through RNA-Seq analysis, focusing on compatibility between species/genotypes and the differentially expressed genes between graft combinations (Rubio et al., 2022; Bantis et al., 2021; Sallaku et al., 2022; Wang et al., 2019; Cookson et al., 2013). In contrast, the phenotypic evidence and the impact on yield and its components were not well documented. These studies showed contrasting results, and the benefit of a given root/shoot combination appears to depend on the specific case, so that no general conclusion can be drawn. In soybean, only one study (Li et al., 2019; see above), using cultivars released in different years to understand the root contribution to breeding, explicitly reported data for both homo- and heterografted plants in the same assay. The authors did not conclude on the benefit of either combination. In the remaining literature, homografts appear only as controls. To the best of our knowledge, the effect of homo- versus heterografting on plant performance has not been specifically addressed in soybean and remains poorly explored in other species. All the mentioned studies grafted plants that differ in their whole genotype rather than in a single gene. The use of transgenic plants in a common Williams 82 background is therefore a promising tool to evaluate root-shoot interactions

Members of the homeodomain-leucine zipper (HD-Zip) family of transcription factors are plant-specific regulators that integrate developmental and environmental signals. These proteins combine a DNA-binding homeodomain with a leucine zipper motif that mediates dimerization, enabling the regulation of gene expression networks involved in growth, hormone signaling, and stress responses (Perotti et al., 2017, 2023). Within this family, the subfamily I members HaHB11 (H11, for the sake of shortness) and HaHB4 (H4), isolated from sunflower, have been described as conferring enhanced yield and abiotic stress tolerance to Arabidopsis plants (Dezar et al. 2005; Cabello et al., 2016; Cabello et al., 2017), and also to crops such as soybean, rice, wheat, and maize(Ribichich et al., 2019; Raineri et al., 2019; Raineri et al., 2022; Raineri et al., 2023; Raineri et al., 202

In soybeans, transgenic expression of *HaHB11* resulted in increased grain number per plant, associated with changes in flowering dynamics, biomass partitioning, and carbon allocation (Raineri et al., 2025). However, since this transcription factor was constitutively expressed, the origin of the observed phenotypic effects, whether driven by the shoot, the root, or their interaction, remains unclear. Moreover, it is unknown at which level (developmental, physiological or molecular), and at which stage of the life cycle, these effects were established. In this context, grafting offers a unique opportunity to dissect the contribution of root and shoot systems to the observed phenotypes.

Here, we use a grafting approach in a uniform soybean genetic background (Williams 82) to evaluate the role of HaHB11 in regulating grain number and yield formation, using also transgenic plants expressing the related transcription factor HaHB4 (which also enhanced grain number; Ribichich et al., 2020) as a second HD-Zip I to test whether the effects are gene-specific rather than a general consequence of expressing a transcription factor of this family. Given the large number of possible graft combinations, the mechanistic analyses focused on the best-performing combination and its relevant controls. By combining transgenic and non-transgenic scions and rootstocks, we integrated field performance with physiological, histological and molecular analyses. We first asked whether the enhanced grain production is primarily controlled by the shoot, the root, or their interaction. We then examined at which level, developmental, physiological or molecular, and at which stage of the life cycle this control was established.

## Results

### 1. Heterografting of transgenic soybeans expressing *HaHB11* and *HaHB4* enhances plant performance across environments

To evaluate the relative contribution of roots and shoots expressing *HaHB11* or *HaHB4* to the increased pod number phenotype, different combinations of rootstocks and scions from Williams 82 control (C), *HaHB4* (H4), and *HaHB11* (H11) soybean lines were generated. Grafting was performed under growth chamber conditions, and plants were subsequently transferred to the field at the V3-V4 stage, and were grown under well-watered conditions in pest- and weed-free soil until physiological maturity (Figure 1).

**Figure 1:**
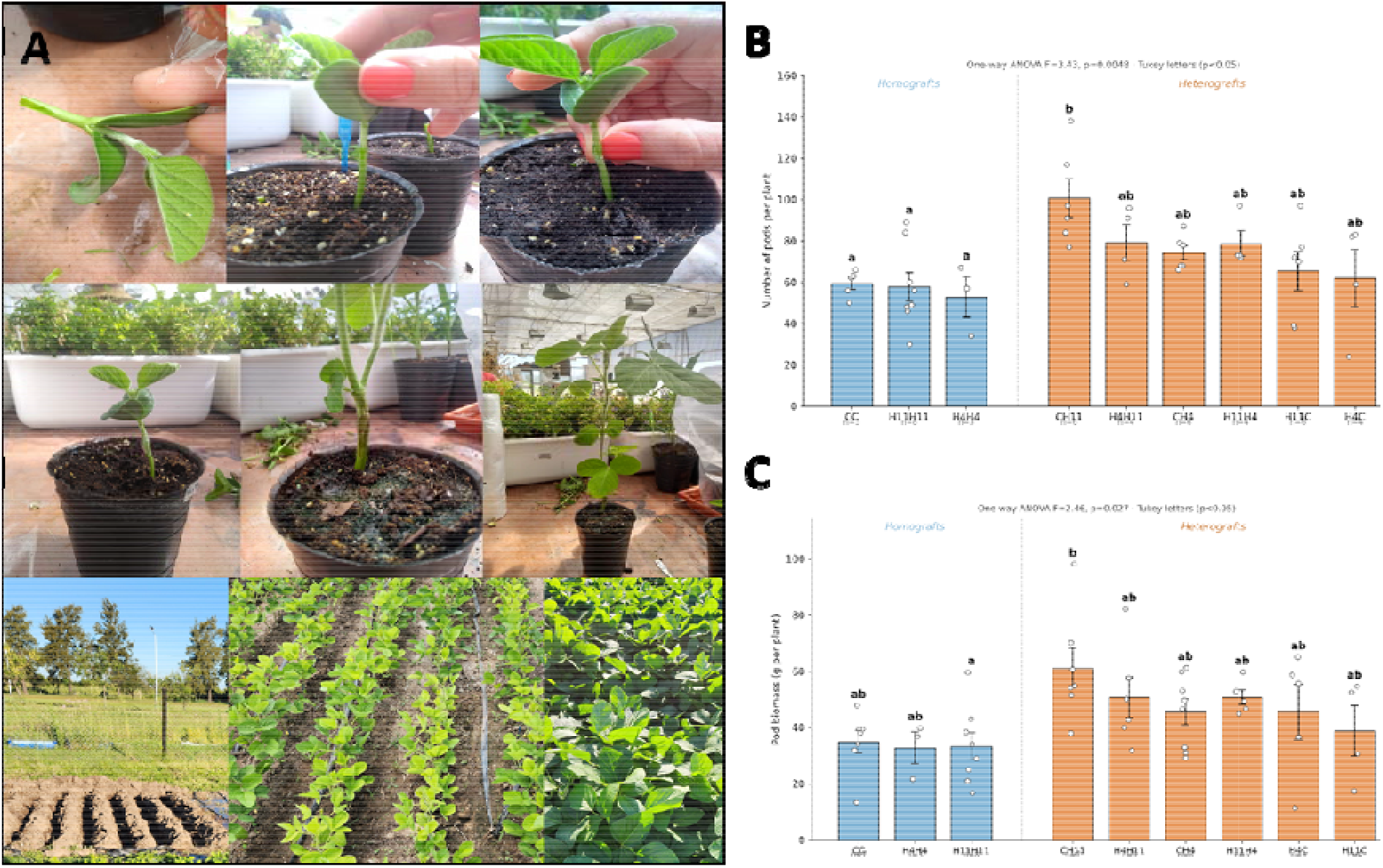
Pod number and pod biomass differences in grafted plants grown in the field depends on the genotype combination. (A) Representative images of the grafting process, recovery, transplanting to the field assay, and crop development. Pod number (B) and pod biomass (C) per plant of the graft combinations. Homografted plants (CC, H11H11, H4H4) are shown in light blue bars and heterografted plants in orange bars (root-scion notation: first letter, rootstock; second letter, scion). Data were analyzed using a one-way ANOVA, followed by Tukey’s HSD post-hoc test; different letters indicate significant differences (P < 0.05).

Unexpectedly, in the first campaign, heterografted plants (CH11, H11C, CH4, H4C; being the first symbol the root donor and the second one the scion donor) tended to show higher pod number and pod biomass than their homografted counterparts (CC, H4H4, H11H11; Figure 1). To further investigate this observation, homo and heterografted plants were evaluated across four environments spanning three growing seasons, including two sowing dates within one season. The joint analysis across environments revealed consistent and significant differences among graft combinations in multiple phenotypic traits. Heterografted plants consistently outperformed homografted ones across environments (Table 1, Supplementary Table S1). Heterografts showed higher number of pods and nodes, stem biomass, total biomass, pod biomass, together with increased stem diameter and higher number of pods/node, relative to homografted controls (Table 1, Supplementary Table S1). This advantage of heterografts over homografts was observed even when both shared the same scion genotype, indicating that the grafting combination itself (beyond the genotype of either organ) modulates plant performance.

**Table 1.** HaHB11 and Williams 82 heterografted plants consistently outperform homografted plants Summary of the comparison of morphological and yield-related traits between homografted and heterografted plants across four environments. For each trait, a linear model was fitted including graft type (homograft vs heterograft) and environment as fixed factors, and both groups were compared through a single linear contrast of estimated marginal means. Full estimated marginal means (±SE) are reported in Supplementary Table S1. Asterisks indicate the significance of the heterograft versus homograft differences. Homografts: CC, H11H11 and H4H4 (same genotype in scion and rootstock); heterografts: all combinations with different scion and rootstock genotypes. *** P < 0.001, ** P < 0.01, * P < 0.05.

|  | Stem diameter | No. of branches | No. Of nodes | No. Of pods | Total biomass | Pods biomass | No. Of pods/node | HI |
| --- | --- | --- | --- | --- | --- | --- | --- | --- |
| Heterograft vs Homograft | * | ** | * | **** | **** | **** | **** | * |

### 2. The scion is the primary driver of yield-related traits in grafts performed with control or transgenic *HaHB4* and *HaHB11* lines

To analyze root-scion interactions, we performed a principal component analysis (PCA) of the yield-related traits evaluated at harvest using rootstock and scion genotypes as grouping factors. PCA analysis supported the association between scion genotype and yield-related traits observed in the phenotypic analyses. PC1 (63.5%) reflected a productivity gradient associated with pod number and biomass-related traits, while PC2 (19.6%) was associated with architectural traits such as branching and individual pod biomass. Figure 2 shows that H4 and, particularly, H11 scions were associated with higher pod number, biomass, and other yield-related traits. Accordingly, heterografted plants carrying H11 or H4 scions (CH4 and CH11) are related to higher pod number and yield than plants with control scions (H4C and H11C; Figure 2).

**Figure 2:**
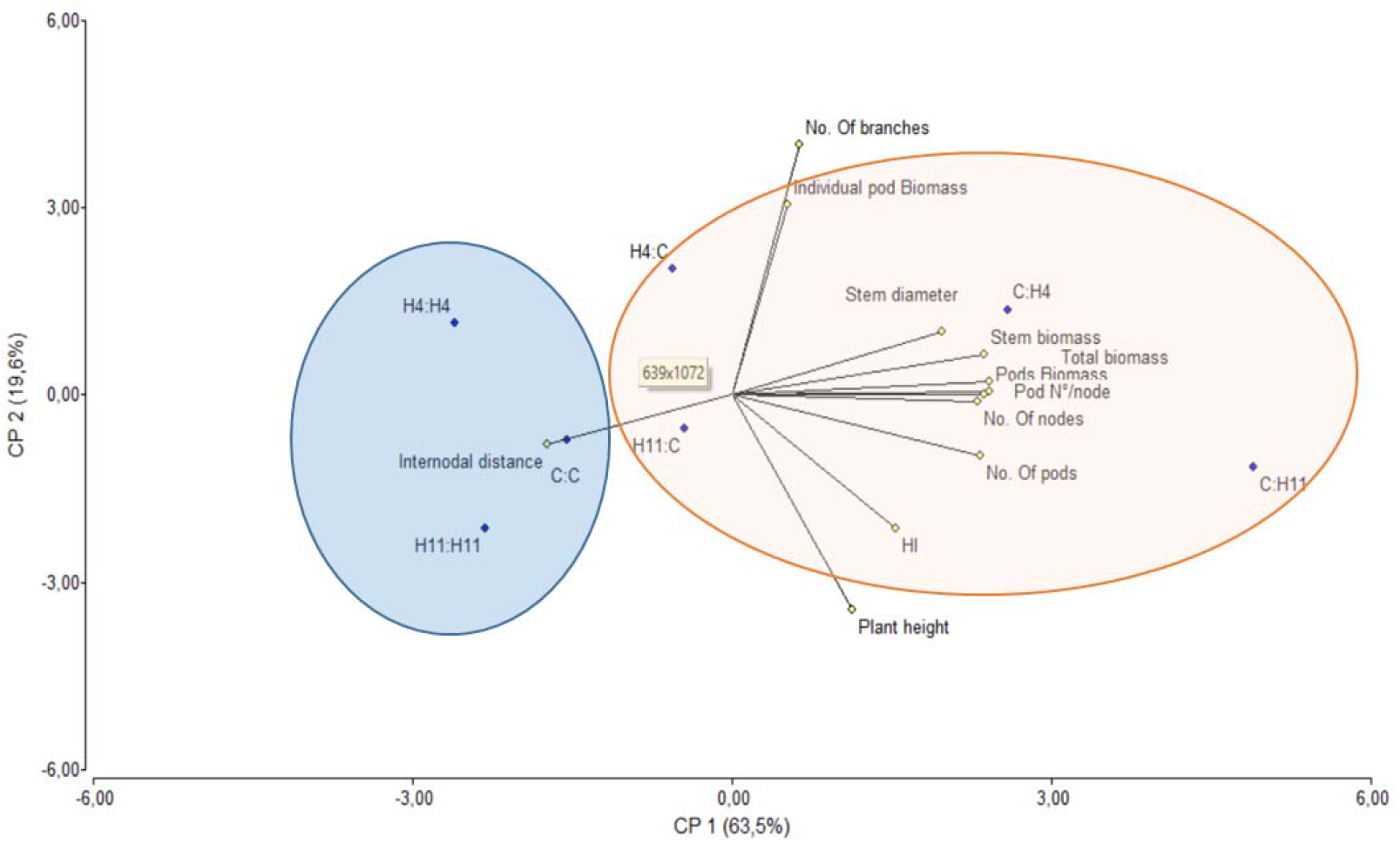
The traits of the combination of control roots and *HaHB11* scions are consistently associated with increased yield. Principal component analysis (PCA) biplot showing relationships among graft combinations and plant traits. Points represent mean values for each rootstock-scion combination. Arrows indicate the contribution of variables to the principal components. Homografted plants (CC; H11H11; H4H4) are grouped in the light blue circle, and heterografted plants in the orange one (root-scion notation: first letter for rootstock, and second letter for scion). Analysis was performed using InfoStat (see Materials and Methods for details on environments included per combination).

Consistently, when individual traits were analyzed using ANOVA, plants grouped by rootstock genotype showed that plants carrying H11 rootstocks had fewer nodes and larger internodal distances compared to the control (C). In contrast, H4 rootstocks had no significant effect on any of the evaluated traits (Table 2, Supplementary Table S2). When plants were grouped by scion genotype, several traits were significantly higher in plants with H11 scions. These traits include plant height, stem diameter and biomass, node and pod number (Table 2, Supplementary Table S2). Plants with H4 scions exhibited a higher harvest index (HI) than plants carrying C scions.

**Table 2.** H11 scions exert stronger effects on plant phenotype than rootstocks. Effects of scion and rootstock genotypes on plant architectural and yield-related traits across all environments and graft combinations. Grafted plants were grouped according to their scion or rootstock genotype. Asterisks indicate the significance of differences between H11 and H4 compared to the control genotype (C). Type II ANOVA was performed, followed by Dunnett-adjusted contrasts of estimated marginal means from linear models including rootstock, scion, and environment as fixed factors. Asterisks indicate significance levels (*** P < 0.001, ** P < 0.01, * P < 0.05), while “–” indicates no significant differences. Green and red shadows denote higher and lower values compared to the control genotype (C). Analyses were performed using all available data, including both homo- and heterografted plants.

| Factor | Genotype | Plant height | Stem diameter | No. of nodes | No. of pods | Stem biomass | HI | Internodal distance |
| --- | --- | --- | --- | --- | --- | --- | --- | --- |
| SCION | H11 | *** | ** | *** | * | * | - | - |
| SCION | H4 | - | - | - | - | - | ** | - |
| ROOT | H11 | - | - | * | - | - | - | * |
| ROOT | H4 | - | - | - | - | - | - | - |

When all combinations were compared (Table 3), an H11 scion reached a comparable high pod number on both a control and an H4 rootstock, but not on its own H11 rootstock, where it dropped to the level of homografts. The same pattern was observed for the H4 scion.

**Table 3:**
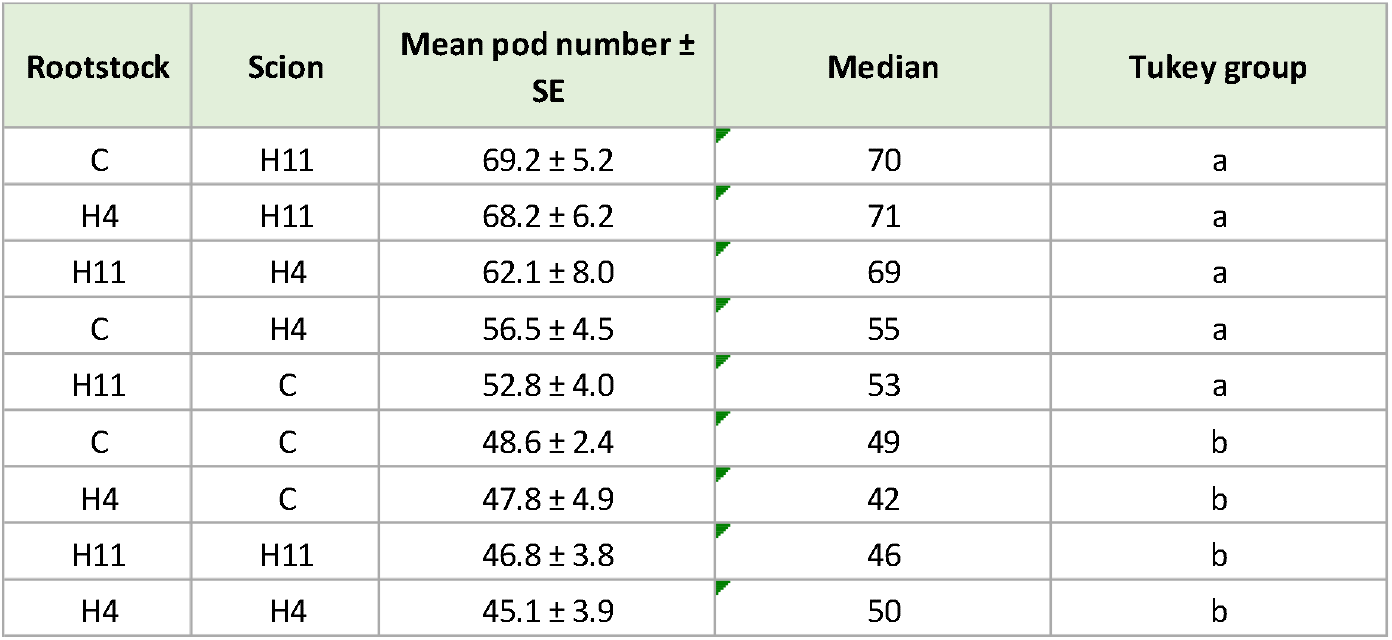
Plants with transgenic H4 and H11 scions exhibit a bigger grain number than those with control scions in heterografted plants. Pod number across all evaluated grafting combinations (Williams 82 background). Plants were pooled across four environments. Combinations were compared by one-way ANOVA with Tukey-adjusted pairwise contrasts. Different letters indicate significant differences (P < 0.05).

These results supported the hypothesis that the yield advantage was associated with combining a given transgenic scion with a rootstock of a different genotype, rather than with the genotype of either organ alone. Besides, the shoot exerts a stronger and more consistent influence on yield-related traits, whereas the rootstock contributes to the variation of specific traits. Together, these findings indicated that both organs regulate plant performance, with a predominant, albeit not limited, role of the shoot in the evaluated conditions, with the rootstock contributing to specific traits rather than to overall yield.

### 3. The scion governs reproductive development, with a minor late-stage rootstock contribution

To examine in which developmental stage each organ exerts its influence, we followed reproductive development over time in plants carrying all combinations of C, H4 and H11 genotypes. Flowering time and development during the reproductive stage were assessed from the emergence of the first bloom (R1) until maturity (R8). Data were analyzed by grouping plants according to either rootstock or scion genotype. The scion genotype is the primary determinant of flowering time, the onset of seed filling and the progression through the reproductive stage (Figure S1, Supplementary Table S3). In contrast, the rootstock genotype affected development only at a late transition: at 112 days after sowing, a significantly lower proportion of plants with H11 rootstocks had reached R7 compared with plants carrying C or H4 rootstocks, which did not differ from each other (Figure S1, Supplementary Table S3). That indicated that the delay in the R6→R7 transition is specific to *HaHB11* expression in the root and not a generic consequence of grafting a transgenic line. Together, these results suggested that the scion governs flowering time and most of reproductive development, whereas the rootstock contributed only at a specific, late stage.

Across the different graft combinations, the scion consistently determined yield-related traits. That prompted a detailed analysis of the best-performing combination.

### 4. Control roots and *HaHB11*-expressing scions grafts are the best-performing combination

CH11 was the combination with the highest mean pod number (Table 3). Based on the presented results and on the phenotypical observations performed, subsequent analyses focused on four representative combinations: CC, H11H11, CH11, and H11C. We selected these grafted combinations to dissect the basis of the CH11 phenotype and the shoot-root interactions.

At physiological maturity, CH11 plants produced more seeds and accumulated higher seed biomass than the other graft combinations (Figure 3A-B). They also exhibited wider stem diameter and a higher number of fertile nodes (Figure 3C-D). CH11 plants showed a tendency toward greater total leaf area, particularly in the lower part of the main stem canopy at R5 (Figure 3E-G). Despite these differences, CH11 plants entered the reproductive phase (R1) synchronously with H11 homografted plants but were delayed compared to plants carrying a control (C) scion (Supplementary Figure S2). The delay in reproductive development of H11 scion plants was maintained until the R4 stage (59 DAS). At 71 DAS, all grafted plants reached the R5 stage (Figure S3).

**Figure 3:**
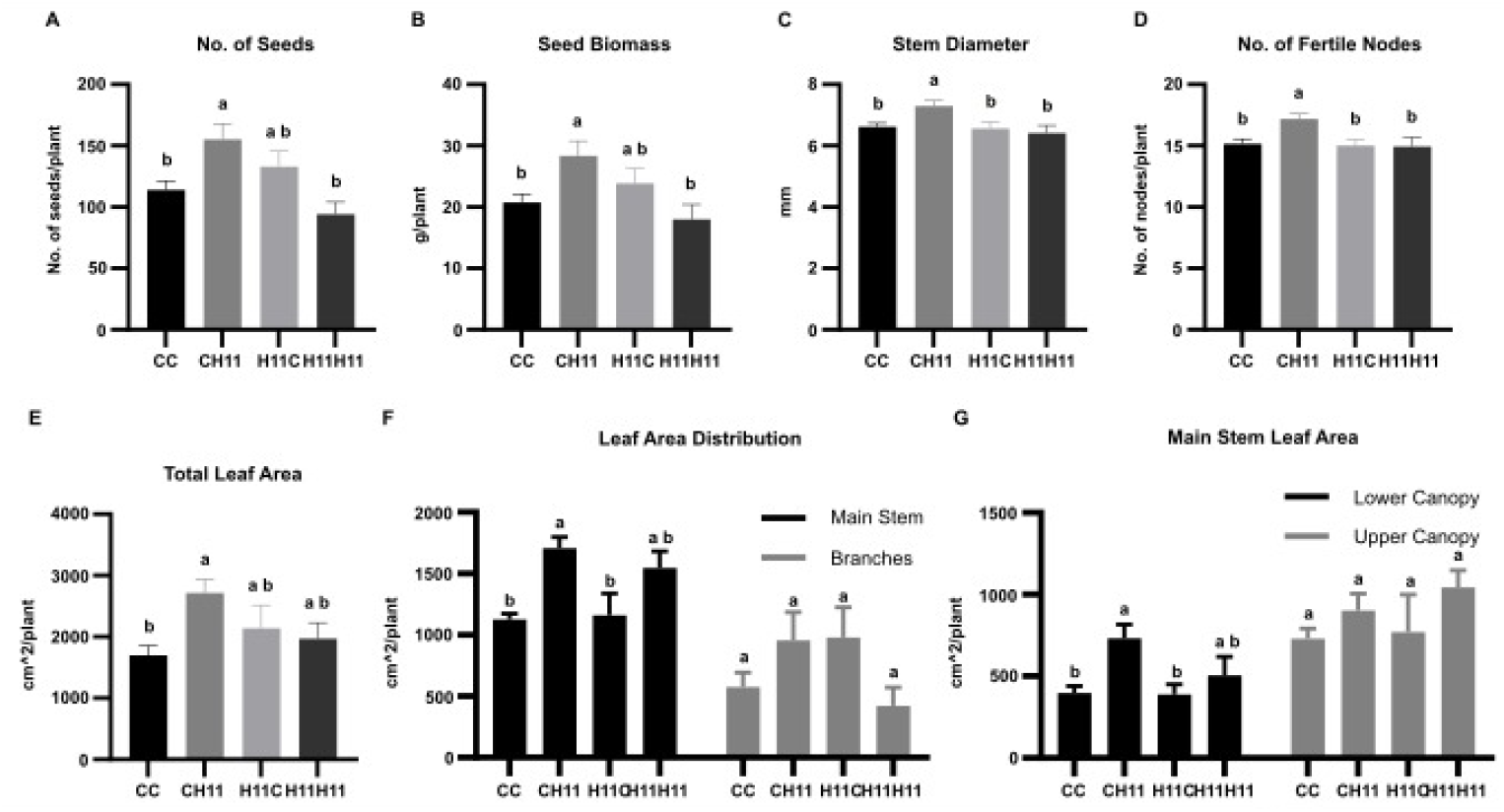
Heterografted CH11 plants exhibit morphological characteristics associated with yield increase. Several traits related to increased yield were assessed for all shoot-root combinations at harvest. (A) Seed number per plant. (B) Seed biomass per plant. (C) Stem diameter. (D) Number of fertile nodes per plant. (E) Total leaf area in R5. (F) Leaf area in stem and branches. (G) Leaf area in the upper and lower parts of the canopy. Data were analyzed using a one-way ANOVA, followed by Tukey’s HSD post-hoc test; different letters indicate significant differences (P < 0.05).

To further evaluate the distribution of pod biomass and individual pod weight throughout the plant, and the possible influence of root proximity on the yield components, the stem was divided into basal, medial, and apical sections. Besides, branches and stems were analyzed separately. The distribution of pod biomass across plant sections showed a trend in plants carrying an H11 scion toward lower allocation to branches and higher allocation to the medial stem; however, these differences were not statistically significant (Supplementary Figure S3). Similarly, individual pod weight is lower in the basal stem section of plants with H11 scions and showed a comparable pattern in branches (Supplementary Figure S3). The proportion of pods containing one, two, three, or four seeds did not differ among graft combinations (Supplementary Figure S3). These results are consistent with the predominant effect of the scion genotype on pod number, as observed in the transversal analysis.

While differences in mature plant architecture and developmental timing contribute to the enhanced performance of CH11 plants, we asked next whether earlier events could underlie this advantage.

### 5. Control roots and *HaHB11*-expressing scion grafts’ advantage may be related to early leaf and stem traits

To determine whether early developmental and physiological differences could contribute to the better performance of CH11 plants, vegetative traits and photosynthetic parameters were evaluated in growth chamber conditions. Stomatal conductance and leaf temperature were measured at the V2-V3 developmental stage. CH11 plants exhibited higher stomatal conductance than the other graft combinations (Figure 4A) and lower leaf temperature (Figure 4B). Heterografted plants showed lower leaf temperature and higher stomatal conductance than homografted plants (Figure 4). To further investigate the basis of increased gas exchange, leaf anatomy was examined. Epidermal peels from the abaxial leaf surface were stained with safranin and used to assess stomatal density. CH11 plants showed significantly higher stomatal density than CC and H11C, while H11H11 plants displayed intermediate values without significant differences from any other combination (Figure 4C-D). No significant differences were observed among combinations in other morphological and physiological traits at V2-V3, including stem length, leaf number, stem diameter, biomass accumulation and partitioning, leaf area, pigment content, and photosynthetic efficiency (Supplementary Table S4).

**Figure 4:**
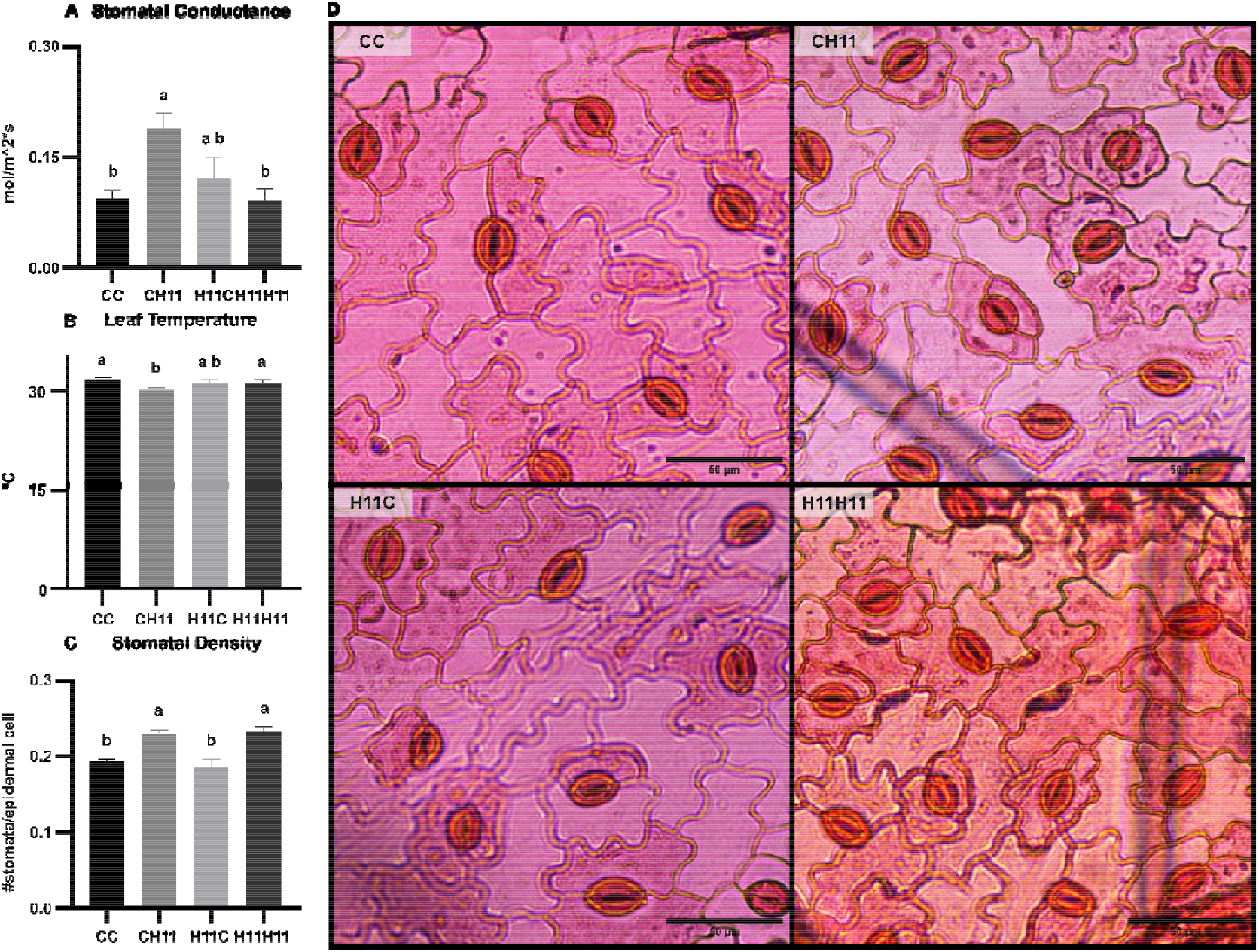
In the V2-V3 early stages of development, CH11 leaves showed higher stomatal conductance. Stomatal conductance (gsw) of the leaf abaxial face (A) and leaf temperature (B) were measured with LICOR 600. (C) Stomatal density expressed as Stomata#/cell#. (D) Illustrative images of abaxial leaf peels for each shoot-root combination. The last expanded leaf was used for analyses. Data were analyzed using a one-way ANOVA. Differences among graft combinations were assessed by one-way ANOVA followed by Tukey’s HSD post-hoc test; different letters indicate significant differences (P < 0.05).

At the vegetative stage, the traits that distinguished CH11 from the other combinations were its higher stomatal conductance and lower leaf temperature, whereas the higher stomatal density were shared with H11H11 and therefore reflected the H11 scion rather than the CH11 combination. These early functional differences in gas exchange, rather than structural ones, may underlie the enhanced performance of CH11 plants. To further investigate the molecular basis of these early differences, we next examined the leaf transcriptome at the same developmental stage.

### 6. The leaf transcriptome is scion-determined and does not single out CH11

To investigate the molecular basis underlying the phenotypic differences observed among graft combinations, an RNA-seq analysis on leaf tissue at the V2 developmental stage was performed, using three biological replicates of CC, CH11, H11C, and H11H11 plants grown in the growth chamber. As expected, plants carrying H11 scions showed high levels of *HaHB11* transcripts (ranging from 31,000 to 101,000 reads). Whereas only background levels (6-118 reads) were detected in plants with control (C) scions, including heterografted plants H11C. This suggests that *HaHB11* transcripts are not detectably transported from roots to shoots in our experimental conditions.

Principal component analysis of the RNA-Seq data showed that the first two components explained 51.9% of the total variance (PC1: 29.6%; PC2: 22.3%). Samples distributed along PC1 according to their scion genotype: CC and H11C replicates grouped on the negative side, while H11H11 and CH11 grouped on the positive side, indicating that scion identity is the main driver of transcriptomic variation in leaves. PC2 captured greater dispersion among heterografted samples compared to homografts (Figure 5A).

**Figure 5:**
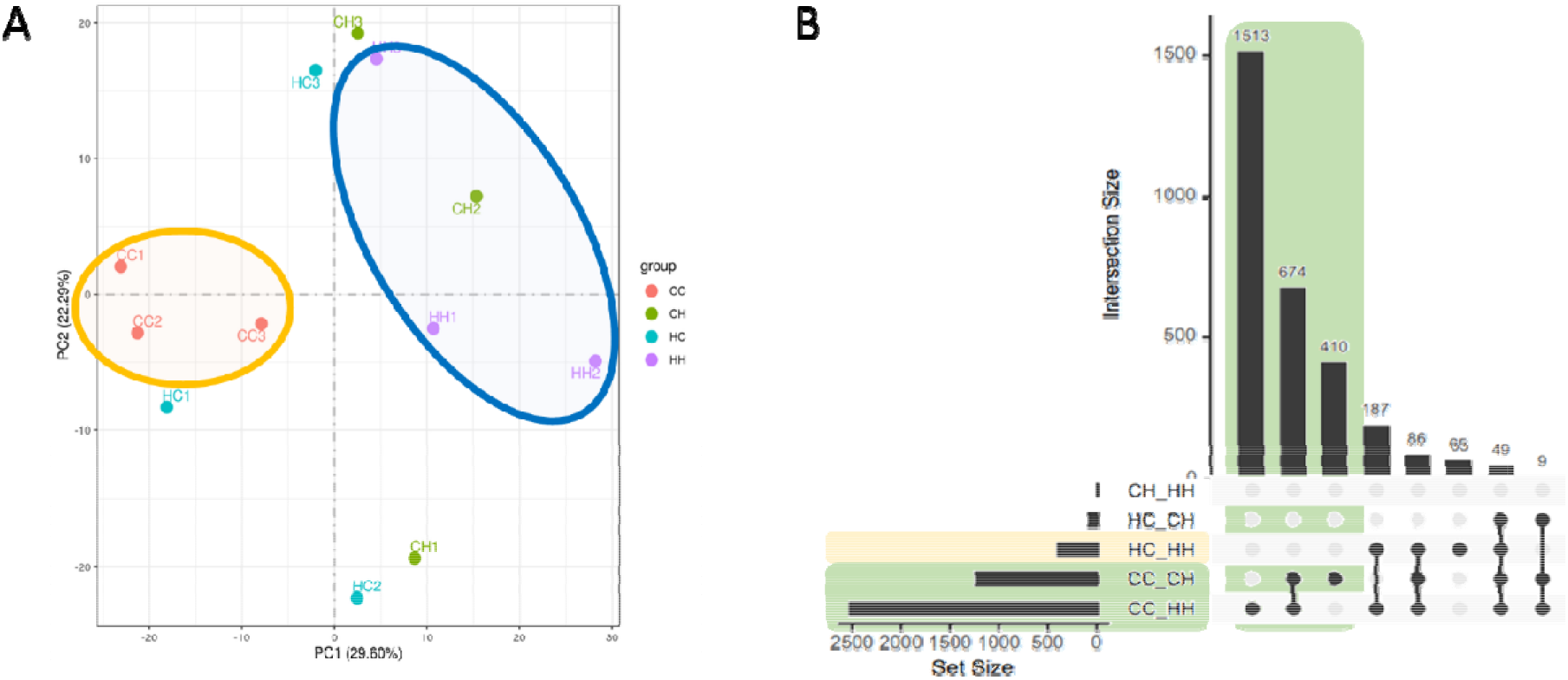
CH11 plants showed 1238 DEG with C homografted, but only four DEG with H11 homografted plants. (A) PCA showing the distribution of the RNA-Seq analysis for biological replicates of each grafted plant’s leaf tissue. CC and H11H11 homografted replicates are shown in orange and violet, respectively. Heterografted H11C and CH11 are in light blue and green. (B) Upsetplot showing the DEG for all shoot-root combinations and the DEG in common between them. CC and H11C plants have no DEG. In green shadow are highlighted the comparisons whose DEGs are related to photosynthesis by the soybean GO analysis.

Pairwise comparisons revealed that the largest number of differentially expressed genes (DEGs) was observed between homografted combinations (CC vs H11H11; 2527 DEGs). Comparisons between CC and CH11 identified 1238 DEGs, approximately half of which overlapped with those detected in CC vs H11H11 (Figure 5B, Supplementary Table S5). In contrast, fewer DEGs were observed between H11H11 and H11C (402 DEGs), with 187 genes shared with the CC vs H11H11 comparison. A substantial proportion of DEGs was shared across these comparisons, indicating the presence of a common transcriptional response associated with H11 scions. In contrast, comparisons between plants sharing the same scion genotype revealed minimal transcriptional differences: only 4 DEGs between H11H11 and CH11, and no DEGs were detected between CC and H11C (Figure 5B, Supplementary Table S5). Heterografted combinations also showed few differences between each other, with only 84 DEGs detected between H11C and CH11. Overall, the number of DEGs followed a gradient consistent with a progressive shift in transcriptomic profiles depending on graft type and scion genotype.

To identify molecular pathways or cellular processes associated with the H11 scion, all DEG sets were analyzed for gene ontology (GO) enrichment in SoyBase and by MapMan analysis using the Arabidopsis homologs of each soybean gene ID. Comparisons between CC and combinations carrying an H11 scion (CC vs CH11 and CC vs H11H11 intersection, 674 genes) showed strong enrichment in photosynthesis-related categories, including photosynthesis, non-photochemical quenching, chlorophyll biosynthesis, photosystem II assembly, and response to light stimulus (Figure 5B; Supplementary Table S6). The MapMan overview revealed consistent transcriptional reconfiguration in light reactions and pigment metabolism between these comparisons (Supplementary Figure S4).

Notably, CH11 plants did not exhibit a unique transcriptional signature in leaf tissue compared to other graft combinations: pairwise comparisons of CH11 against H11H11 and against H11C identified very few DEGs (4 and 85, respectively), with no significantly enriched GO categories in the comparisons. Thus, although the H11 scion imposes a clear transcriptional program on the leaf, this program does not distinguish CH11 from H11H11, indicating that the superior performance of CH11 is not explained by a distinct leaf transcriptome.

Since CH11 did not exhibit a unique leaf transcriptomic signature, we next explored whether metabolic reconfiguration could contribute to its enhanced reproductive phenotype. For this purpose, GC-MS-based metabolite profiling was performed on leaf tissue from the four focal graft combinations at solar noon and before lights-off.

### 7. The leaf metabolome shows scion and rootstock components but no CH11 specific signature

To assess whether the enhanced grain number of CH11 plants was correlated to metabolic rearrangements in the leaf, we profiled polar metabolites by GC-MS across the four grafting combinations. Samples were taken at V2- V3 plants grown in a growth chamber at two contrasting times of day, Zeitgeber Time 8 (ZT8, i.e., 8 hours after lights-on) at solar noon and ZT16, end of day. Diurnal accumulation of the main photosynthetic sugars (sucrose and glucose, higher at ZT16 than at ZT8; Supplementary Tables S7-S8) confirmed that the profiles captured physiologically coherent dynamics. The relative abundance of all detected metabolites is shown in Figure 6. While no graft combination showed a globally distinct metabolic profile, several metabolites varied in a genotype-specific manner. Consistently, principal component analysis did not resolve a clear separation between graft combinations at either time (Supplementary Figure S5). Grouping plants as homo- versus heterografted yielded no metabolite that survived multiple-testing correction. Thus, the grafting combinations did not differ in their global leaf metabolome but in a restricted set of metabolites, which we resolved into scion- and rootstock-associated components.

**Figure 6.**
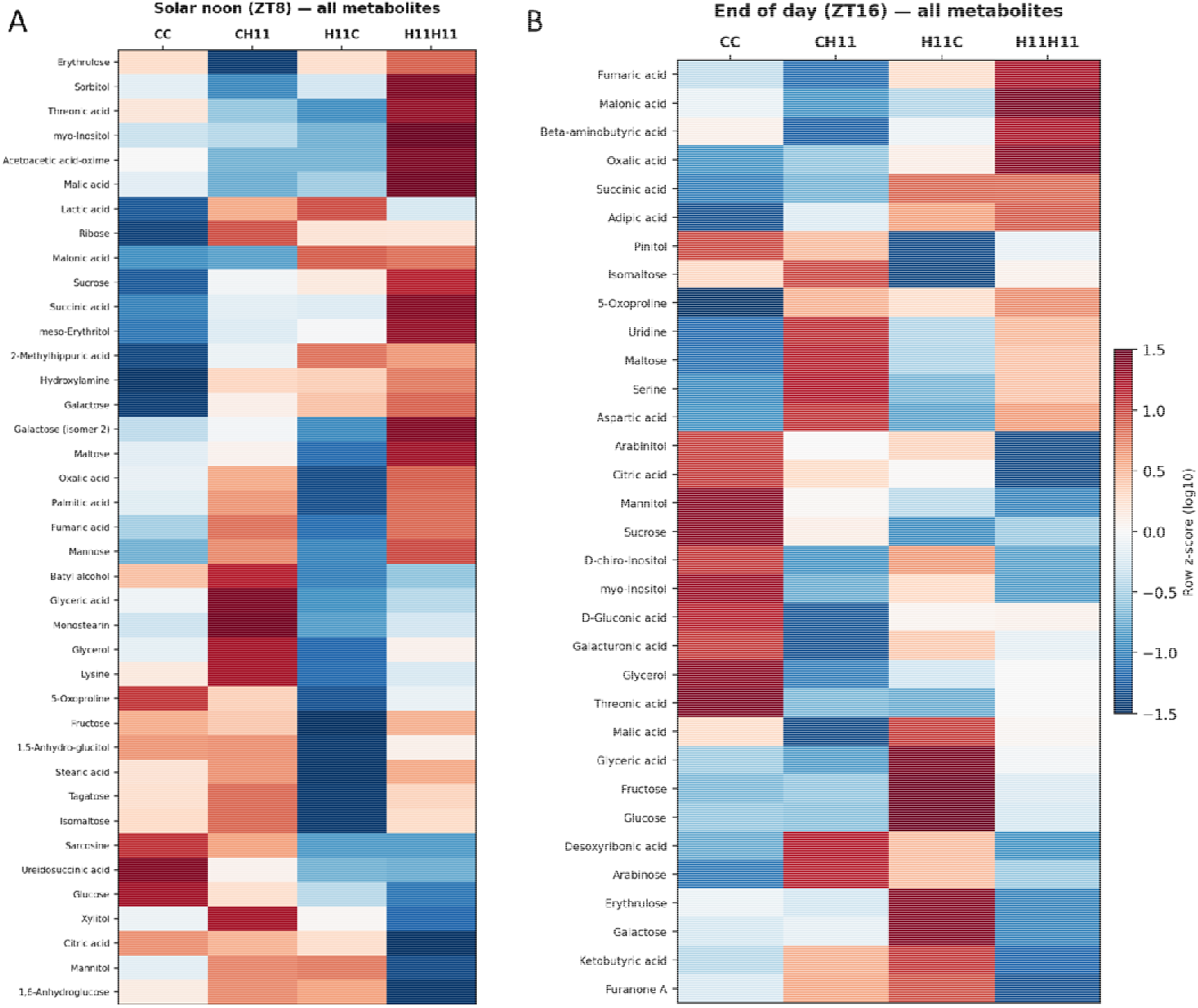
Leaf metabolomes are largely conserved across grafting combinations, with few metabolite-specific differences. Heatmaps of polar metabolites measured by GC-MS in leaves of the four focal grafting combinations (CC, CH11, H11C, H11H11) at (A) solar noon (ZT8, eight hours after lights-on) and (B) end of day (ZT16, sixteen hours after lights-on). Values are row-wise z-scores of log₁₀-transformed mean abundances per grafting combination (n = 3 biological replicates at ZT8; n = 4-5 at ZT16; each replicate is a pool of leaves from three plants). Colors are comparable across genotypes within a metabolite (row) but not between metabolites. Red indicates higher and blue lower abundance relative to the mean of the four genotype means. The scale was capped at ±1.5. Rows are ordered by hierarchical clustering (Euclidean distance, average linkage). Columns are shown in fixed order.

To test whether the scion or the roots have main effects on the leaf metabolome, we analyzed data grouped by tissue genotype. The scion genotype accounted for the larger share of changes between combinations, consistent with its predominant role in the phenotypic and transcriptomic data (Supplementary Figure S6, Supplementary Tables S9-S10). Plants bearing an H11 scion showed a consistent, non-significant tendency toward higher maltose at both times of the day, whereas the remaining scion-associated metabolites differed between samplings: mannose and fumaric acid were elevated at ZT8 (both q < 0.05), while at ZT16, myo-inositol and malic acid were reduced (the only two metabolites significant after correction at this time).

A second, smaller component of the signature tracked the rootstock genotype. Succinic acid was reproducibly higher in plants carrying an H11 rootstock at both times of day, and at ZT16,these plants also showed higher fumaric acid and lower citric acid (Supplementary Tables S9-S10). Metabolite set enrichment analysis (MSEA, KEGG) of the scion- and rootstock-associated metabolites revealed significant pathway-level structure in a two contrasts: Scion- affected metabolites at ZT16 were enriched in Glyoxylate cycle and dicarboxylate metabolism (FDR= 0.013) and rootstock-affected metabolites at ZT16 were enriched in the TCA cycle (FDR = 2.6 × 10⁻⁵; Supplementary Table S11), pointing to a concurrent scion-rootstock influence on central carbon metabolism at the end of the day. This rootstock-associated metabolic component contrasts with the leaf transcriptome, which was determined almost exclusively by the scion genotype. Notably, and as observed for the transcriptome, CH11 did not display a distinctive metabolic signature: no metabolite set it apart from the other combinations, and in particular, it did not differ from H11H11. Thus, neither the leaf transcriptome nor the leaf metabolome singles out CH11, pointing to its early architectural and physiological traits as the basis of its advantage.

Collectively, H11 overexpression drives modest but reproducible shifts in leaf primary metabolism, centred on the TCA cycle and soluble sugar pools, arising predominantly from the scion with a smaller yet consistent rootstock contribution. Although individual changes were small, their consistency across two independent samplings supports their biological relevance.

## Discussion

### Heterografted plants outperformed homografted plants

We observed that heterografted plants performed better than homografted plants, considering several physiological traits during the vegetative and reproductive developmental stages (Table 1, Figure 1). In contrast, leaf transcriptomic and metabolomic profiles did not reveal differences specifically distinguishing homo- from heterografted plants (Figures 5 and 6). Notably, there are no previous documents reporting beneficial effects on soybean yield of heterografting. Li et al. (2019) grafted two record-yield cultivars (L14 and Z35) as scions onto rootstocks of historical cultivars, and their data showed lower seed number, seed biomass and harvest index in the homografted controls compared to the heterografts. Unfortunately, the authors did not test or discuss this interesting experiment. When these authors analyzed the reciprocal combinations, using the record-yield cultivars as rootstocks, they observed that other parameters, such as seed weight, harvest index, and yield were higher in heterografts than in homografts (Li et al., 2017b). It is important to note that these heterografts combined historical and record-yield genotypes; hence, the effect of the graft combination itself cannot be separated from that of the high-yielding genotype. Our experimental design used isogenic lines differing in a single heterologous gene, allowing us to isolate the graft-combination effect, suggesting that the observed yield advantage arose from combining a scion with a rootstock of a different genotype, rather than from the yield potential of each genotype.

Notably, H11 homografted plants did not differ in grain number or yield from control homografted plants, even though non-grafted H11 plants outyielded controls (Raineri et al., 2025), suggesting that the homografting procedure itself attenuated the grain number advantage conferred by HaHB11.

### The scion is the primary organ determining pod number

The analysis of grouping plants by scion or rootstock genotype identified the scion as the primary organ determining pod number under control conditions in the Williams 82 background, whereas the rootstock played a minor role (Table 2). Similar assays aimed to dissect the influence of the root and the shoot on different phenotypic traits in soybean, including photosynthesis and yield components (Du et al., 2019; Li et al., 2017), nodulation (Streeter, 1974; Delves et al., 1986; He et al., 2021; Wang et al., 2021), and flowering (Przepiorkowski et al., 2003; Cober & Curtis, 2003). Although the authors identified one organ as predominant over the phenotype, the non-dominant organ generally exerted minor but non-negligible effects on the evaluated traits. None of these studies used transgenic plants. Cober and Curtis (2003) evaluated early- and late-flowering soybean genotypes by grafting and showed that, in early-flowering genotypes, flowering is mainly controlled by the shoot, whereas in late-flowering genotypes the rootstock genotype allowed plants to reach R1 earlier. While the rootstock had a minor overall influence, its contribution was neither random nor diffuse but temporally and functionally circumscribed. Such observations were similar to our results, in which the scion was the main organ determining flowering transition and grain number; the roots also contributed with a minor effect. Evaluating a very different trait, fibrous roots, Pantalone et al. (1999) worked with grafted soybean plants of the line PI4 (with fibrous roots) and plants of the line Lee74. They concluded that the root itself regulated the fibrous-like trait, albeit they also reported an effect on seed weight associated with root genotype. In contrast with our results, these authors did not report a scion effect and attributed all changes to the root genotype. So, the relative contribution of root or shoot to the phenotype may be genotype- and trait-dependent.

### The leaf transcriptome and metabolome reflect genotype rather than graft type

The leaf transcriptomic analysis of graft combinations was consistent with the results presented above: the scion genotype governed the transcriptomic profile (Figure 5). Unlike our findings, He et al. (2021) used two soybean genotypes contrasting in nodulation (NN1138-2, with normal nodulation and T3791, a non-nodulating mutant) and reported that the scion and the rootstock genotypes reciprocally shaped gene expression in both leaves and roots. To our knowledge, only one study has analyzed the reciprocal grafting transcriptome of near-isogenic soybean lines: Kohlhase et al. (2025) examined shoot and root transcriptomes of plants differing in tolerance to iron-deficiency chlorosis. This work analyzed plants under acute iron deficiency and reported that within 30–120 min of stress the leaf transcriptome shifted from being determined by the leaf’s own genotype to being governed by the rootstock. Together, these contrasting results suggested that the organ governing the leaf transcriptome is not fixed, but condition-dependent. Under the non-stress conditions evaluated in that work, the scion was the determinant, whereas the rootstock can take over this control under acute abiotic stress, or even the root-shoot interaction played a part in a complex trait such as nodulation. The hierarchy of long-distance transcriptional control may thus be reconfigured by the environment.

mannose and fumaric acid were elevated at ZT8 (both q < 0.05), while at ZT16, myo-inositol and malic acid

Leaf metabolomic analysis also showed a predominant scion component, identifying individual metabolites sourviving FDR test only in this tissue. Although, both organs left metabolic signatures associated with organic acid metabolism (glyoxylate in the scion and TCA in the rootstock; Supplementary Tables S9-S11). Whether this contributes to the grain-number phenotype remains to be elucidated. To our knowledge, there are no other works on grafted soybean that include leaf tissue metabolomics assays. In line with the rootstock component we detected, in other systems the root genotype and the scion-root interplay (including their compatibility) shaped the metabolic profile of the plant aerial part. Specifically, the citrus fruit and the scion of salt-stressed tomato showed rootstock-dependent changes in TCA-cycle intermediates (Tietel et al., 2020; Song et al., 2022).

### Some of the HaHB11 characteristics are common to different species

#### The flowering delay and the increased leaf area generated by HaHB11 can be atributed to the scion

In all transgenic species tested, *HaHB11* overexpression impacted their architecture and development. Moreover, Arabidopsis, rice, maize, and soybean H11 plants showed a delay in the onset of flowering. In the same way, grafted H11 scion plants have a delay in reaching the R1 stage, compared to C scion plants (Supplementary Figure S1-S2 and Table S3). As in H11 scion-grafted plants, in Arabidopsis, rice and maize H11 plants exhibited larger leaf area than controls (Cabello et al., 2016; Raineri et al., 2022, Figure 3), pointing to a conserved phenotype conferred by the transgene, affecting plant development in normal conditions.

#### H11 scions outperformed other combinations only when heterografted

Among all graft combinations, CH11 was the highest-yielding, and its increased seed yield was mainly explained by seed number. The advantage of CH11 plants was related to early physiological traits (including higher stomatal conductance and lower leaf temperature; Figure 4) rather than structural traits shared with H11 homografts (higher stomatal density) or distinct molecular signatures. The H11 scion reached a high pod number when combined with a different rootstock (CH11 or H4-rooted plants), but not on its own H11 rootstock (Table 3), whose yield fell to control levels. Thus, the advantage conferred by HaHB11 in the scion was expressed only in a heterografted context, indicating that the graft combination, rather than the scion genotype alone, determines whether the HaHB11 benefits are expressed (Figure 7).

**Figure 7.**
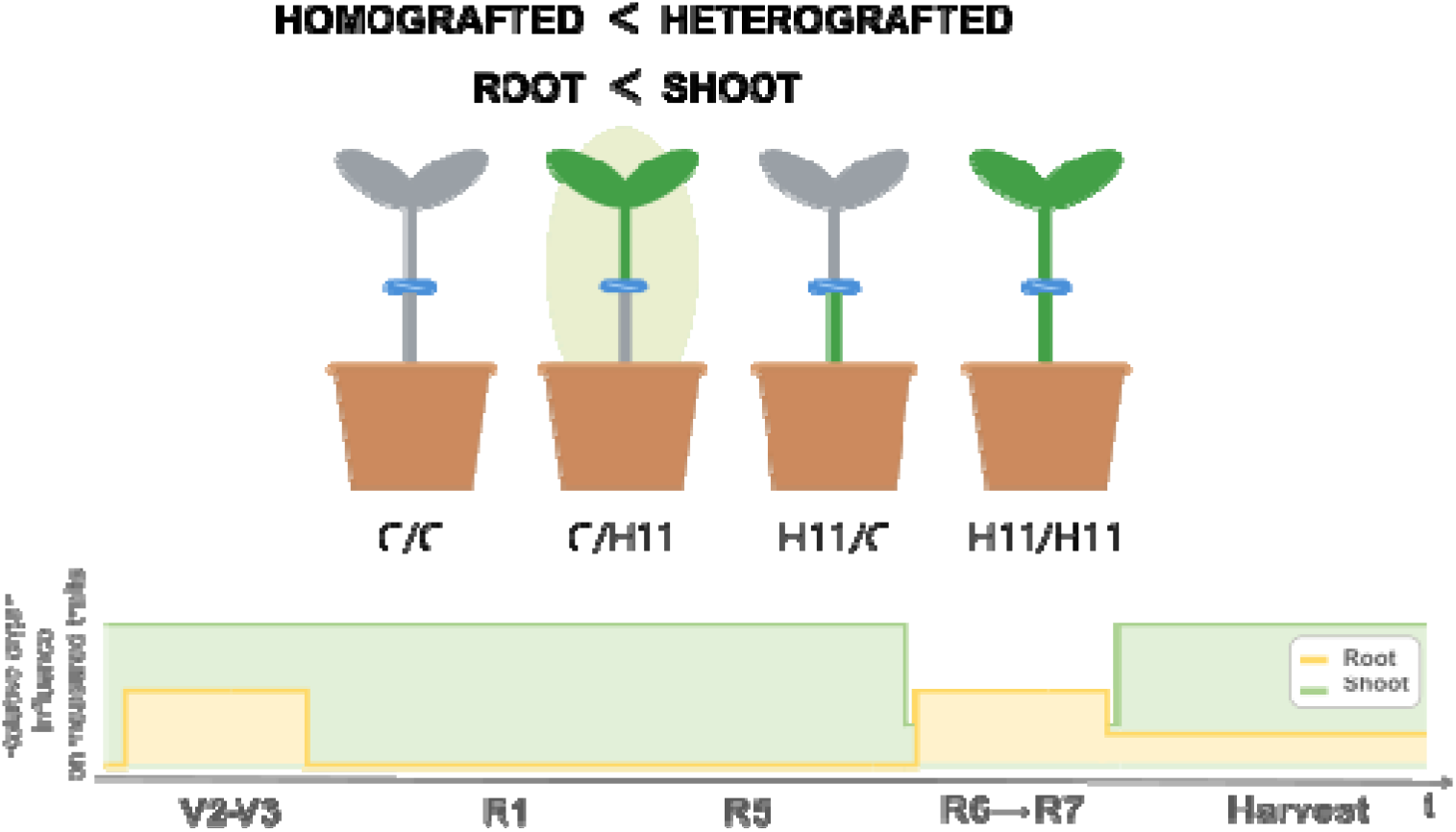
Heterografting with an H11 scion enhances soybean performance, with organ influence shifting across development. (Top) Reciprocal grafts between Williams 82 control (C) and *HaHB11*-overexpressing (H11) lines. Green tissue denotes the H11 genotype; grey/brown tissue denotes the control. (Bottom) Schematic synthesis of the relative influence of the root (yellow) and shoot (green) on the traits measured at each developmental stage. This diagram is a qualitative integration of the results and not a quantitative estimate of organ contribution: the readouts differ between stages (leaf transcriptome, metabolome and gas-exchange traits at V2-V3; flowering and reproductive progression at R1-R5; developmental-transition timing at R6→R7; yield components at harvest).

Further work may be necessary to reveal whether the graft-combination effect is HD-Zip specific and/or depends on transcriptional reprogramming. For this purpose, it will be necessary to use isogenic lines expressing transcription factors from other families or genes playing different roles, and extend the analysis to stress conditions and root tissues.

The strength of this work lies in the reciprocal grafting of isogenic lines differing in a single gene (*HaHB11*) in a common Williams 82 background, which allowed us to separate the contribution of each organ from that of the graft combination itself. Our results indicated that, under non-stress conditions, grain number in soybean grafts is governed mainly by the scion, whereas the graft combination determines whether the HaHB11 benefit is expressed. To our knowledge, this is one of the first studies to evaluate yield and yield-related traits in field-grown soybean grafts combined with leaf transcriptomic and metabolomic analyses of growth-chamber-grown plants.

## Materials and Methhods

### Plant material and growth conditions

Seeds were germinated and grafting was performed in growth chamber conditions following Pantalone et al. (1999). For homografted plants, the scion and the roots came from different plants with the same genotype. Due to variation in post-grafting survival rates in the growth chamber, the number of plants available for field transplantation varied across assays, ranging from 4 to 25 plants per graft combination. Surviving plants were transplanted to the field at the V3 stage, exhibiting a post-transplantation survival rate exceeding 99%.For V2–V3 phenotyping, grafted plants were grown on chamber, or transferred to the greenhouse and maintained until the V2–V3 developmental stage, when phenotypic and physiological evaluations were performed. Greenhouse temperature ranged from 28 to 36 °C and relative humidity from 70 to 90 % during the experiment. Growth chamber temperature was set at 28°C, and relative humidity from 60 to 90%.

### Field experimental design

Field assays were carried out at the IAL site (31°38’S, 60°40’W ,Santa Fe, Argentina) on a sandy soil of 2 m depth with low water-holding capacity and an organic ’A’ horizon of 15 cm. Grafted plants were evaluated across four environments spanning three growing seasons (2021–2022, 2023–2024, and 2024–2025). Within the 2021–2022 season, two sowing dates were used (December 2^nd^ and December 17^th^), constituting two independent environments (Env. 1 and Env. 2, respectively). The 2023–2024 season (Env. 3, sown on December 13^th^) and the 2024–2025 season (Env. 4, sown on December 10^th^) each constituted a single environment. Graft combinations involving the Williams 82 background, being C the wild-type plants and H4 and H11, transgenic plants expressing the *HaHB4* and *HaHB11* sunflower transcription factors, respectively. CC, CH11, H11C, H11H11, CH4, H4C, and H4H4 grafted plants were evaluated in Env. 1, 2, and 3. Env. 4 included only the four focal combinations (CC, CH11, H11C, and H11H11). In Env. 4, a higher number of plants per combination was achieved due to the reduced number of graft combinations evaluated. Within each environment, grafted plants were arranged in randomized blocks of two to three plants, with at least three blocks per environment; the position of every plant in the plot was recorded. For statistical analyses, environment (Env.) was included as a fixed factor in the models to account for environmental variation.

Seeds were inoculated with *Bradyrhizobium japonicum* before sowing. Plots were drip-irrigated throughout the cycle to maintain the uppermost 1 m soil layer at field capacity and were fertilized with P (130 kg ha⁻¹ at sowing). Parcels were kept free of weeds, insects, and diseases. Daily maximum temperature (°C) and incident solar radiation (MJ m⁻² day⁻¹) were recorded from a nearby meteorological station. All experiments were conducting according to the regulatory organisms rules.

### Evaluation of crop phenotype

Crop phenotype evaluation was performed essentially as described in Raineri et al., 2025. Reproductive development was scored at four time points after sowing (40, 79, 112 and 121 days after sowing, DAS). At each date, the reproductive stage of each plant was recorded according to Fehr and Caviness (1977), and plants were classified according to whether they had reached or surpassed a given reproductive stage (≥R1, ≥R6, ≥R7, ≥R8).

### Stomatal conductance, leaf temperature, and PSII quantum efficiency

Stomatal conductance (gsw), leaf temperature, and PSII quantum efficiency (ΦPSII) were taken with a LI-600 Porometer/Fluorometer (LI-COR Biosciences, Lincoln, NE, USA) on the abaxial surface of the youngest fully expanded leaf at V2–V3. Measurements were taken at solar noon under photosynthetically active radiation (PAR) ranging from 400 to 600 µmol m⁻² s⁻¹. ΦPSII was calculated for at least four plants per graft combination, by the instrument using a multiphase saturating flash according to Loriaux et al. (2013).

### Stomatal density

Epidermal peels of the abaxial surface of the youngest fully expanded leaf at V2–V3 were obtained, stained with safranin, and mounted on glass slides. Between 7 and 10 microscopic fields per leaf were photographed using an Eclipse E200 microscope (Nikon, Tokyo, Japan) coupled to a Nikon Coolpix L810 camera. Stomata and total epidermal cells were counted manually in each field. Stomatal density was expressed as the ratio of the number of stomata to the total number of epidermal cells. One leaf per plant and at least four plants per graft combination, and seven to ten fields of each sample were analyzed.

### Pigment quantification

Chlorophyll *a*, chlorophyll *b*, and carotenoid contents were quantified spectrophotometrically. One leaf disk (4 mm radius) per plant was placed in 1.5 mL of 80 % acetone (acetone:milliQ water, v/v) and incubated overnight in darkness with gentle agitation. Absorbance was recorded at 663, 646, and 470 nm against a reagent blank. Chlorophyll and carotenoid concentrations were calculated according to Porra et al. (1989) and Lichtenthaler (1987), respectively. Pigment content was expressed per unit leaf area (µg/cm²).

### Anatomical analysis of stem vascular tissue

Hand-cut transverse sections of the epicotyl were obtained at V2–V3 and stained with safranin/fast green. Sections were photographed using an Eclipse E200 microscope (Nikon, Tokyo, Japan) coupled to a Nikon Coolpix L810 camera. The relative area of secondary xylem was quantified using ImageJ (NIH, Bethesda, MD, USA) and expressed as the proportion of the total cross-sectional area of the stem.

## Statistical analyses

For the transversal evaluation (Tables 1-2, Supplementary Tables S1-S2), statistical analyses were performed using R (version 4.5.3; R Core Team, 2026) on data pooled across environments, with environment (Env.) included as a fixed factor to account for environmental variation. All linear models used Type II sums of squares (car package), and estimated marginal means and contrasts were obtained with the emmeans package. Significance was considered at P < 0.05, and 0.05 ≤ P < 0.10 was interpreted as a marginal trend.

To assess the overall effect of graft type (Supplementary Table S1), all plants were grouped as homografted (same genotype in scion and rootstock: CC, H11H11, and H4H4) or heterografted (combinations with different scion and rootstock genotypes). For each trait, a linear model was fitted including graft type and environment as fixed factors, and the two groups were compared through a single linear contrast of estimated marginal means (heterograft vs homograft).

To evaluate the effect of each graft component (Supplementary Tables S2), linear models were fitted for each trait including rootstock genotype (Root), scion genotype (Shoot) and environment as fixed factors. Differences relative to the control genotype (C) were assessed using estimated marginal means with Dunnett-adjusted contrasts.

For Figure 2 Principal Component Analysis (PCA) was performed in InfoStat (Di Rienzo et al., 2020) on the mean values of each rootstock-scion combination for the yield-related and architectural traits. The analysis was based on the correlation matrix (standardized variables). H11-containing combinations were evaluated across four environments and H4-containing combinations across three environments.

For Supplementary Table S3, the progression of reproductive development (from R1 to R8) was analyzed by grouping plants according to their scion or their rootstock genotype. At each sampling date (40, 79, 112 and 121 days after sowing, DAS), the proportion of plants that had reached or surpassed a given reproductive stage was compared among genotypes using χ² contingency tests. P-values were corrected for multiple testing with the Benjamini–Hochberg false discovery rate (FDR) procedure, applied separately within each factor (scion or rootstock) across sampling dates. When the global test was significant (q < 0.05), pairwise comparisons among genotypes were performed using Fisher’s exact tests with FDR correction. These analyses were carried out in Python (SciPy).

To directly compare graft combinations (Table 3, Figures 3-4, Supplementary Figures S2-3, Supplementary Table S4), each trait was analyzed separately using a one-way analysis of variance (ANOVA), with the rootstock–scion combination treated as a single factor. Post hoc comparisons were performed using Tukey-adjusted pairwise contrasts based on estimated marginal means. Statistical groupings were summarized using compact letter displays.

For GC-MS statistical analysis, metabolite levels (ribitol-normalized peak areas per mg fresh weight) were log_10_-transformed to better approximate normality. The two sampling times (ZT8 and ZT16) were analyzed separately, as they corresponded to independent batches of plants. For each metabolite, the effects of scion genotype, rootstock genotype, and their interaction were assessed by two-way ANOVA (type II sums of squares). The effect of each graft component was further evaluated by grouping plants by scion (C scion: CC + H11C; H11 scion: CH11 + H11H11) or by rootstock (C rootstock: CC + CH11; H11 rootstock: H11C + H11H11) and comparing groups with Welch’s t-tests; an analogous contrast grouped plants as homo- (CC + H11H11) versus hetero-grafted (CH11 + H11C). P-values were corrected for multiple testing within each sampling time using the Benjamini–Hochberg false discovery rate (FDR) procedure (Benjamini & Hochberg, 1995). These univariate analyses were performed in Python 3.12 (SciPy v1.17, NumPy, pandas).

Pairwise comparisons among the four grafting combinations (Tukey’s HSD post-hoc test) and PCA were carried out in MetaboAnalyst 5.0 (Pang et al., 2021) on log10-transformed data; for PCA, data were additionally autoscaled (unit variance). Metabolite set enrichment analysis (MSEA) was performed in MetaboAnalyst on the metabolites associated with each graft component (scion or rootstock, p < 0.10) at each sampling time, using the KEGG pathway-associated metabolite set library. As MetaboAnalyst does not provide a plant-specific pathway library, human KEGG metabolite sets were used; central carbon metabolism pathways (e.g. the TCA cycle) are conserved across kingdoms, and enrichment was interpreted accordingly. Pathway enrichment was considered significant at FDR < 0.05.

Heatmaps were generated in Python (Matplotlib) and display all metabolites with complete data across the four grafting combinations (i.e. detected in at least one replicate of each genotype), excluding the internal standard (ribitol) and derived metabolite ratios. For each metabolite, log_10_-transformed values were averaged per grafting combination, and values shown are row-wise z-scores of these genotype means, so colors are comparable across genotypes within a metabolite (row) but not between metabolites.

## RNA-seq analysis

Three biological replicates from leaf of V2 each combination of grafted plants grown in growth chamber (16:8 light:dark phoyoperiod, 28°) were taken at ZT8. RNA integrity was assessed using the Bioanalyzer 2100 system (Agilent Technologies, CA, USA). Messenger RNA was purified from total RNA using poly-T oligo-attached magnetic beads. After fragmentation, the first strand cDNA was synthesized using random hexamer primers. Then the second strand cDNA was synthesized using dUTP, instead of dTTP. The directional library was ready after end repair, A-tailing, adapter ligation, size selection, amplification, and purification. The library was checked with Qubit and real-time PCR for quantification and bioanalyzer for size distribution detection. Sequencing was performed on a Illumina NovaSeq X Plus instrument (Illumina Inc., CA, USA), obtaining 150 bp paired-end reads and a sequencing depth between 45 and 78 million reads. The bioinformatics pipeline started with a quality check using FastQC (ver. 0.12.1, Andrews, 2010) and summarized with MultiQC (ver. 1.21, Ewels et al., 2016). Reads were then trimmed and filtered with trimmomatic (ver. 0.39, Bolger et al., 2014), aligned to the Gmax_508_Wm82.a4.v1 genome, obtained from Phytozome (Goodstein et al., 2012), using STAR (ver. 2.7.11b, Dobin et al., 2013), using gene annotation from the same source. Counts per gene were determined with featureCounts (ver. 2.0.6, Liao et al., 2013) and used to determine differentially expressed genes with the DESeq2 package (ver. 1.47.5, Love et al., 2014) in R (R Core Team, 2025), using a cutoff of 0.05 for the adjusted p-value.

## GC-MS-based metabolomic analysis

Polar metabolites were extracted from leaf tissue following the protocol of Lisec et al. (2006). Soybean plants of the four focal grafting combinations (CC, CH11, H11C, and H11H11) were grown in culture chamber under long photoperiod (16:8, Light:dark), 28°C and sampled at the V2-V3 stage at two zeitgeber times (ZT), with lights on at 6:00 h: ZT8 (solar noon) and ZT16 (end of the day). Sampling was performed on independent batches of plants (n = 3 biological replicates at ZT8, and n=4-5 at ZT16). Each biological replicate consisted of a pool of leaves from three plants, taking three central portions of the leaflets from the youngest fully expanded leaf. Tissue was immediately frozen in liquid nitrogen and stored at −80 °C until processing. Between 30 and 50 mg of fresh tissue were used per sample, weighed accurately for subsequent normalization. As internal standard, 30 µL of a freshly prepared aqueous ribitol solution (0.2 mg/mL) were added to each sample before drying.

Derivatization was performed in two steps. First, a freshly prepared methoxyamine hydrochloride solution (20 mg/mL in anhydrous pyridine) was added (40 µL per sample) and samples were incubated at 37 °C with shaking (1000 rpm) for 2 h. Second, 70 µL of MSTFA (N-methyl-N-(trimethylsilyl)trifluoroacetamide) were added, and samples were incubated again at 37 °C with shaking (1000 rpm) for 30 min. Ribitol and methoxyamine blanks were prepared as controls. Derivatized extracts (70 µL) were transferred to GC vials and analyzed on a Nexis GC-2030 gas chromatograph coupled to a GCMS-TQ8050 NX triple quadrupole mass spectrometer (Shimadzu, Kyoto, Japan), equipped with an SH-I-5MS column. Chromatographic conditions were: injection temperature 250 °C, split injection mode (split ratio 10), helium as carrier gas (53.1 kPa, column flow 1 mL/min), oven program 60 °C (1 min), 15 °C/min ramp to 300 °C, held for 5 min. Ion source and interface temperatures were 230 °C and 250 °C, respectively. Solvent cut time was 6 min and acquisition mode was SCAN (m/z 60–800). Chromatographic data were processed with the GC-MS Insight Software Package for TQ Series (Shimadzu), and metabolites were identified by comparison against the Smart Metabolites Database (Shimadzu). Peak areas were normalized to the area of the ribitol internal standard in each sample and then divided by the exact fresh tissue mass (mg).

Metabolites detected in fewer than 50 % of all samples within a time point were considered low-quality features and excluded from downstream analysis. Within the retained set, missing values in individual samples (i.e. metabolites detected in most but not all replicates) were handled by each test using available data; no value imputation was performed.

## Supporting information

Supplementary Tabes

## References

1. Andrews, S., 2010. FastQC: a quality control tool for high throughput sequence data. Babraham Bioinformatics, Babraham Institute, Cambridge, UK. https://www.bioinformatics.babraham.ac.uk/projects/fastqc/

2. Augstein, F., Melnyk, C.W., 2025. Modern and historical uses of plant grafting to engineer development, stress tolerance, chimeras, and hybrids. Plant J. 121, e70057.

3. Bantis, F., Tsiolas, G., Mouchtaropoulou, E., Tsompanoglou, I., Polidoros, A.N., Argiriou, A., Koukounaras, A., 2021. Comparative transcriptome analysis in homo- and hetero-grafted cucurbit seedlings. Front. Plant Sci. 12, 691069.

4. Benjamini, Y., Hochberg, Y., 1995. Controlling the false discovery rate: a practical and powerful approach to multiple testing. J. R. Stat. Soc. Series B Methodol. 57, 289–300.

5. Bolger, A.M., Lohse, M., Usadel, B., 2014. Trimmomatic: a flexible trimmer for Illumina sequence data. Bioinformatics 30, 2114–2120.

6. Cabello, J.V., Giacomelli, J.I., Piattoni, C.V., Iglesias, A.A., Chan, R.L., 2016. The sunflower transcription factor HaHB11 improves yield, biomass and tolerance to flooding in transgenic Arabidopsis plants. J. Biotechnol. 222, 73–83.

7. Cabello, J.V., Giacomelli, J.I., Gómez, M.C., Chan, R.L., 2017. The sunflower transcription factor HaHB11 confers tolerance to water deficit and salinity to transgenic Arabidopsis and alfalfa plants. J. Biotechnol. 257, 35–46.

8. Cao, X., Wu, T., Sun, S., Wu, C., Wang, C., Jiang, B., Tao, J., Yao, W., Hou, W., Yang, W., Siddique, K.H.M., Han, T., 2019. Evaluation by grafting technique of changes in the contribution of root-to-shoot development and biomass production in soybean (Glycine max) cultivars released from 1929 to 2006 in China. Crop Pasture Sci. 70, 585–594.

9. Cober, E.R., Curtis, D.F., 2003. Both promoters and inhibitors affected flowering time in grafted soybean flowering-time isolines. Crop Sci. 43, 886–891.

10. Cookson, S.J., Clemente Moreno, M.J., Hevin, C., Nyamba Mendome, L.Z., Delrot, S., Trossat-Magnin, C., Ollat, N., 2013. Graft union formation in grapevine induces transcriptional changes related to cell wall modification, wounding, hormone signalling, and secondary metabolism. J. Exp. Bot. 64, 2997–3008.

11. Cookson, S.J., Clemente Moreno, M.J., Hevin, C., Nyamba Mendome, L.Z., Delrot, S., Magnin, N., Trossat-Magnin, C., Ollat, N., 2014. Heterografting with nonself rootstocks induces genes involved in stress responses at the graft interface when compared with autografted controls. J. Exp. Bot. 65, 2473–2481.

12. Delves, A.C., Mathews, A., Day, D.A., Carter, A.S., Carroll, B.J., Gresshoff, P.M., 1986. Regulation of the soybean-Rhizobium nodule symbiosis by shoot and root factors. Plant Physiol. 82, 588–590.

13. Dezar, C.A., Gago, G.M., González, D.H., Chan, R.L., 2005. Hahb-4, a sunflower homeobox-leucine zipper gene, is a developmental regulator and confers drought tolerance to Arabidopsis thaliana plants. Transgenic Res. 14, 429–440.

14. Di Rienzo, J.A., Casanoves, F., Balzarini, M.G., Gonzalez, L., Tablada, M., Robledo, C.W., 2020. InfoStat versión 2020. Grupo InfoStat, FCA, Universidad Nacional de Córdoba, Córdoba, Argentina. http://www.infostat.com.ar

15. Dobin, A., Davis, C.A., Schlesinger, F., Drenkow, J., Zaleski, C., Jha, S., Batut, P., Chaisson, M., Gingeras, T.R., 2013. STAR: ultrafast universal RNA-seq aligner. Bioinformatics 29, 15–21.

16. Du, Y., Zhao, Q., Li, S., Yao, X., Xie, F., Zhao, M., 2019. Shoot/root interactions affect soybean photosynthetic traits and yield formation: a case study of grafting with record-yield cultivars. Front. Plant Sci. 10, 445.

17. Ewels, P., Magnusson, M., Lundin, S., Käller, M., 2016. MultiQC: summarize analysis results for multiple tools and samples in a single report. Bioinformatics 32, 3047–3048.

18. Fang, C., Dong, L., Zhou, J., Lu, S., Liu, B., 2026. Toward a Green Revolution in soybean: the role of ultra-high-density planting. J. Integr. Plant Biol. 68, 297–301.

19. Fehr, W.R., Caviness, C.E., 1977. Stages of Soybean Development. Special Report 80. Iowa State University Cooperative Extension Service, Ames, IA.

20. Fuglie, K., 2016. The growing role of the private sector in agricultural research and development world-wide. Glob. Food Sec. 10, 29–38.

21. Goodstein, D.M., Shu, S., Howson, R., Neupane, R., Hayes, R.D., Fazo, J., Mitros, T., Dirks, W., Hellsten, U., Putnam, N., Rokhsar, D.S., 2012. Phytozome: a comparative platform for green plant genomics. Nucleic Acids Res. 40, D1178–D1186.

22. He, Q., Xiang, S., Wang, W., Shu, Y., Li, Z., Wang, S., Chen, L., Yang, X., Zhao, T., 2021. Transcriptomic and photosynthetic responses to grafting of the Nod1 gene in nodulated and non-nodulated soybeans. G3 (Bethesda) 11, jkab209.

23. Jin, J., Liu, X., Wang, G., Mi, L., Shen, Z., Chen, X., Herbert, S.J., 2010. Agronomic and physiological contributions to the yield improvement of soybean cultivars released from 1950 to 2006 in Northeast China. Field Crops Res. 115, 116–123.

24. Koester, R.P., Skoneczka, J.A., Cary, T.R., Diers, B.W., Ainsworth, E.A., 2014. Historical gains in soybean (Glycine max Merr.) seed yield are driven by linear increases in light interception, energy conversion, and partitioning efficiencies. J. Exp. Bot. 65, 3311–3321.

25. Koester, R.P., Nohl, B.M., Diers, B.W., Ainsworth, E.A., 2016. Has photosynthetic capacity increased with 80 years of soybean breeding? An examination of historical soybean cultivars. Plant Cell Environ. 39, 1058–1067.

26. Kohlhase, D.R., O’Rourke, J.A., Graham, M.A., 2025. RNA-seq of grafted near-isogenic soybean (Glycine max) lines reveals root genotype drives shoot responses to iron deficiency chlorosis. Plant Stress 15, 100717.

27. Lam, H.-M., Xu, X., Liu, X., Sun, W., Chen, H., Wong, F.-L., et al., 2010. Resequencing of 31 wild and cultivated soybean genomes identifies patterns of genetic diversity and selection. Nat. Genet. 42, 1053–1059. *[verificar ajuste temático — foco genómico, ver mi nota]*

28. Lee, J.M., Kubota, C., Tsao, S.J., Bie, Z., Hoyos Echevarria, P., Morra, L., Oda, M., 2010. Current status of vegetable grafting: diffusion, grafting techniques, automation. Sci. Hortic. 127, 93–105.

29. Li, S., Teng, F., Rao, D., Yao, X., Zhang, H., Wang, H., Song, S., St Martin, S.K., Xie, F., 2017a. Agronomic traits of soybean cultivars released in different decades after grafting record-yield cultivar as rootstock. Plant Breed. 136, 133–138.

30. Li, S., Teng, F., Rao, D., Zhang, H., Wang, H., Yao, X., Yu, C., Li, C., Zhao, M., St Martin, S.K., Xie, F., 2017b. Photosynthesis of soybean cultivars released in different decades after grafting onto record-yield cultivars as rootstocks. Photosynthetica 55, 579–587.

31. Li, S., Wang, W., Cao, Y., Wang, C., Yan, C., Dong, L., Wu, L., Xie, F., Song, S., 2019. How root traits would be affected by soybean yield improvement? An examination of historical cultivars grafted with record-yield cultivar scion. Plant Soil 439, 19–30.

32. Li, S., Cao, Y., Wang, C., Sun, X., Wang, W., Song, S., 2021. Contribution of different genotypic roots to drought resistance in soybean by a grafting experiment. Plant Prod. Sci. 24, 317–325.

33. Liao, Y., Smyth, G.K., Shi, W., 2014. featureCounts: an efficient general purpose program for assigning sequence reads to genomic features. Bioinformatics 30, 923–930.

34. Lichtenthaler, H.K., 1987. Chlorophylls and carotenoids: pigments of photosynthetic biomembranes. Methods Enzymol. 148, 350–382.

35. Lisec, J., Schauer, N., Kopka, J., Willmitzer, L., Fernie, A.R., 2006. Gas chromatography mass spectrometry-based metabolite profiling in plants. Nat. Protoc. 1, 387–396.

36. Loriaux, S.D., Avenson, T.J., Welles, J.M., McDermitt, D.K., Eckles, R.D., Riensche, B., Genty, B., 2013. Closing in on maximum yield of chlorophyll fluorescence using a single multiphase flash of sub-saturating intensity. Plant Cell Environ. 36, 1755–1770.

37. Loupit, G., Brocard, L., Ollat, N., Cookson, S.J., 2023. Grafting in plants: recent discoveries and new applications. J. Exp. Bot. 74, 2433–2447.

38. Love, M.I., Huber, W., Anders, S., 2014. Moderated estimation of fold change and dispersion for RNA-seq data with DESeq2. Genome Biol. 15, 550.

39. Mudge, K., Janick, J., Scofield, S., Goldschmidt, E.E., 2009. A history of grafting. Hortic. Rev. 35, 437–493.

40. Pang, Z., Chong, J., Zhou, G., de Lima Morais, D.A., Chang, L., Barrette, M., Gauthier, C., Jacques, P.-É., Li, S., Xia, J., 2021. MetaboAnalyst 5.0: narrowing the gap between raw spectra and functional insights. Nucleic Acids Res. 49, W388–W396.

41. Pantalone, V.R., Rebetzke, G.J., Burton, J.W., Carter, T.E., Israel, D.W., 1999. Soybean PI 416937 root system contributes to biomass accumulation in reciprocal grafts. Agron. J. 91, 840–844.

42. Passioura, J.B., 1983. Roots and drought resistance. Agric. Water Manag. 7, 265–280.

43. Perotti, M.F., Ribone, P.A., Chan, R.L., 2017. Plant transcription factors from the homeodomain-leucine zipper family I. Role in development and stress responses. IUBMB Life 69, 280–289.

44. Perotti, M.F., Arce, A.L., Chan, R.L., 2021. The underground life of homeodomain-leucine zipper transcription factors. J. Exp. Bot. 72, 4005–4021.

45. Porra, R.J., Thompson, W.A., Kriedemann, P.E., 1989. Determination of accurate extinction coefficients and simultaneous equations for assaying chlorophylls a and b extracted with four different solvents: verification of the concentration of chlorophyll standards by atomic absorption spectroscopy. Biochim. Biophys. Acta Bioenerg. 975, 384–394.

46. Przepiorkowski, T., St Martin, S.K., 2003. The effect of grafting on the flowering of near-isogenic lines of soybean. Crop Sci. 43, 1760–1763.

47. Raineri, J., Campi, M., Chan, R.L., Otegui, M.E., 2019. Maize expressing the sunflower transcription factor HaHB11 has improved productivity in controlled and field conditions. Plant Sci. 287, 110185.

48. Raineri, J., Caraballo, L.N., Rigalli, N., Portapila, M., Otegui, M.E., Chan, R.L., 2022. Expressing the sunflower transcription factor HaHB11 in maize improves waterlogging and defoliation tolerance. Plant Physiol. 189, 230–247.

49. Raineri, J., Caraballo, L.N., Gómez, M., Chan, R.L., 2023. The transcription factor HaHB11 boosts grain set and yield in rice plants, allowing them to approach their ideal phenotype. Biomolecules 13, 826.

50. Raineri, J., Montero Bulacio, E., Campi, M., Portapila, M., Otegui, M.E., Chan, R.L., 2025. The sunflower transcription factor HaHB11 increases soybean grain number and heat tolerance across multi-season field trials. J. Exp. Bot. 76, 5037–5055.

51. Ribichich, K.F., Chiozza, M., Ávalos-Britez, S., Cabello, J.V., Arce, A.L., Watson, G., Arias, C., Portapila, M., Trucco, F., Otegui, M.E., Chan, R.L., 2020. Successful field performance in warm and dry environments of soybean expressing the sunflower transcription factor HB4. J. Exp. Bot. 71, 3142–3156.

52. Rogers, E.D., Benfey, P.N., 2015. Regulation of plant root system architecture: implications for crop advancement. Curr. Opin. Biotechnol. 32, 93–98.

53. Rubio, B., Stammitti, L., Cookson, S.J., Teyssier, E., Gallusci, P., 2022. Small RNA populations reflect the complex dialogue established between heterograft partners in grapevine. Hortic. Res. 9, uhab067.

54. Sallaku, G., Rewald, B., Sandén, H., Balliu, A., 2022. Scions impact biomass allocation and root enzymatic activity of rootstocks in grafted melon and watermelon plants. Front. Plant Sci. 13, 949086.

55. Song, C., Acuña, T., Adler-Agmon, M., Rachmilevitch, S., Barak, S., Fait, A., 2022. Leveraging a graft collection to develop metabolome-based trait prediction for the selection of tomato rootstocks with enhanced salt tolerance. Hortic. Res. 9, uhac061.

56. Streeter, J.G., 1974. Growth of two soybean shoots on a single root: effect on nitrogen and dry matter accumulation by shoots and on the rate of nitrogen fixation by nodulated roots. J. Exp. Bot. 25, 189–198.

57. Tietel, Z., Srivastava, S., Fait, A., Tel-Zur, N., Carmi, N., Raveh, E., 2020. Impact of scion/rootstock reciprocal effects on metabolomics of fruit juice and phloem sap in grafted Citrus reticulata. PLoS One 15, e0227192.

58. Tollenaar, M., Aguilera, A., 1992. Radiation use efficiency of an old and a new maize hybrid. Agron. J. 84, 536–541.

59. Wang, H., Zhou, P., Zhu, W., Wang, F., 2019. De novo comparative transcriptome analysis of genes differentially expressed in the scion of homografted and heterografted tomato seedlings. Sci. Rep. 9, 20240.

60. Wang, T., Guo, J., Peng, Y., Lyu, X., Liu, B., Sun, S., Wang, X., 2021. Light-induced mobile factors from shoots regulate rhizobium-triggered soybean root nodulation. Science 374, 65–71.

