## Supplementary material for "Isogenic reciprocal grafts with transgenic HaHB11 plants dissect shoot and root contributions to yield in field-grown soybean: a multi-omic study"

**Supplementary Figures and tables**

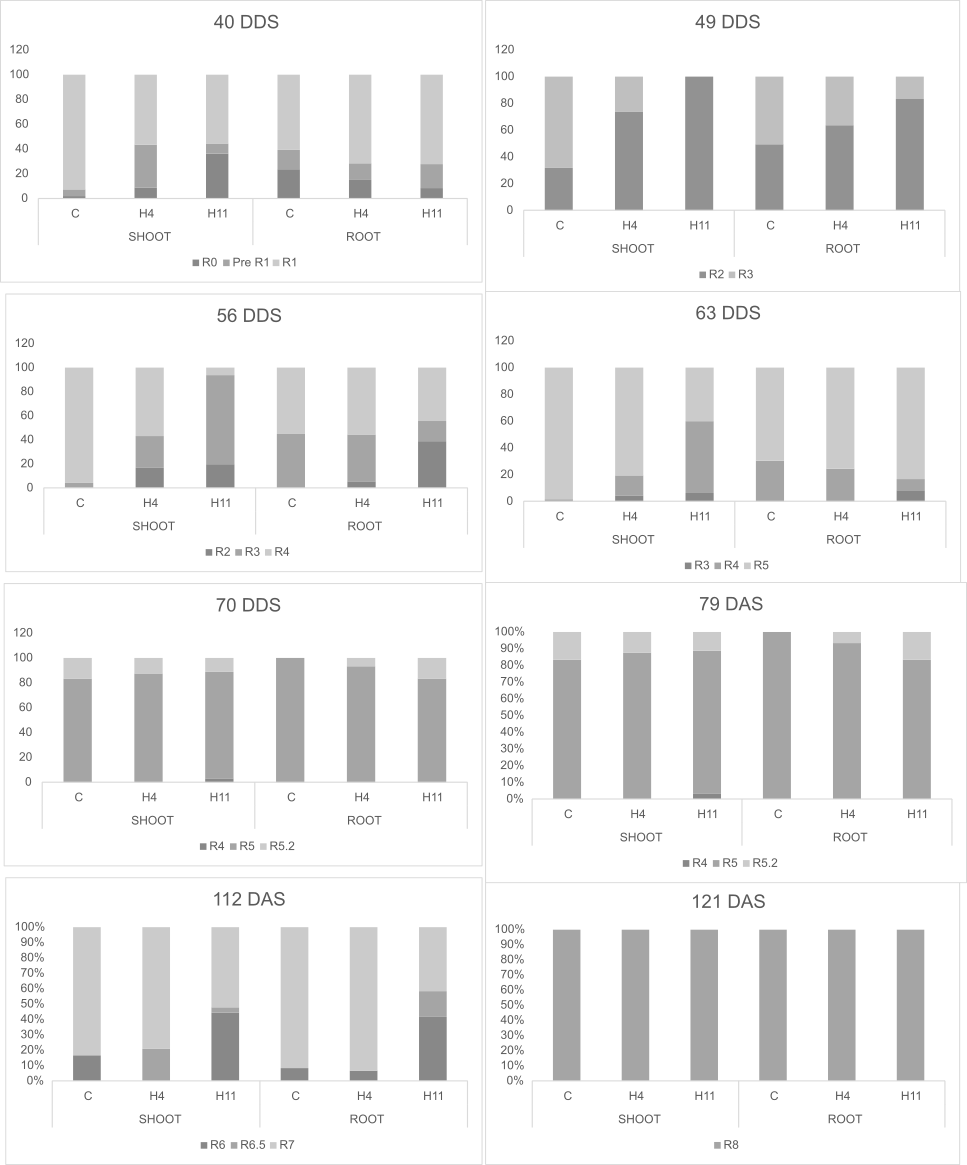

**Supplementary Figure S1. Reproductive development of grafted plants grouped by scion or rootstock genotype**

Grafted soybean plants were grouped according to their scion or rootstock genotype: control (C), *HaHB11*-expressing (H11) and *HaHB4* (H4). At each sampling date (40–121 days after sowing, DAS), the proportion of plants at each reproductive stage (R0–R8, color-coded) was scored within each group. Reproductive development was followed for every individual plant from emergence through flowering (R1) to maturity (R8). Bars show the percentage of plants at each stage. Statistical comparisons of the proportion of plants reaching key reproductive stages are reported in Supplementary Table S3. DAS: days after sowing.

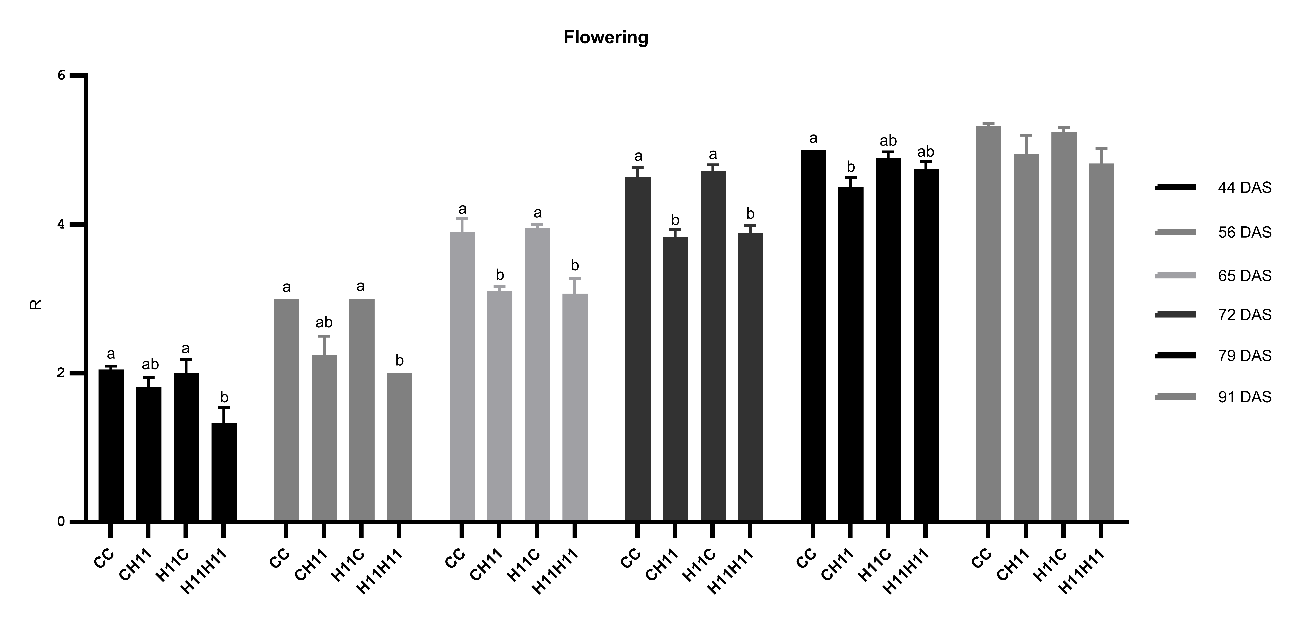

**Supplementary Figure S2. The phenology of the grafted plants does not differ between genotype combinations**

Phenology from flowering (R1) to beginning of grain filling (R5). DAS: days after sowing.

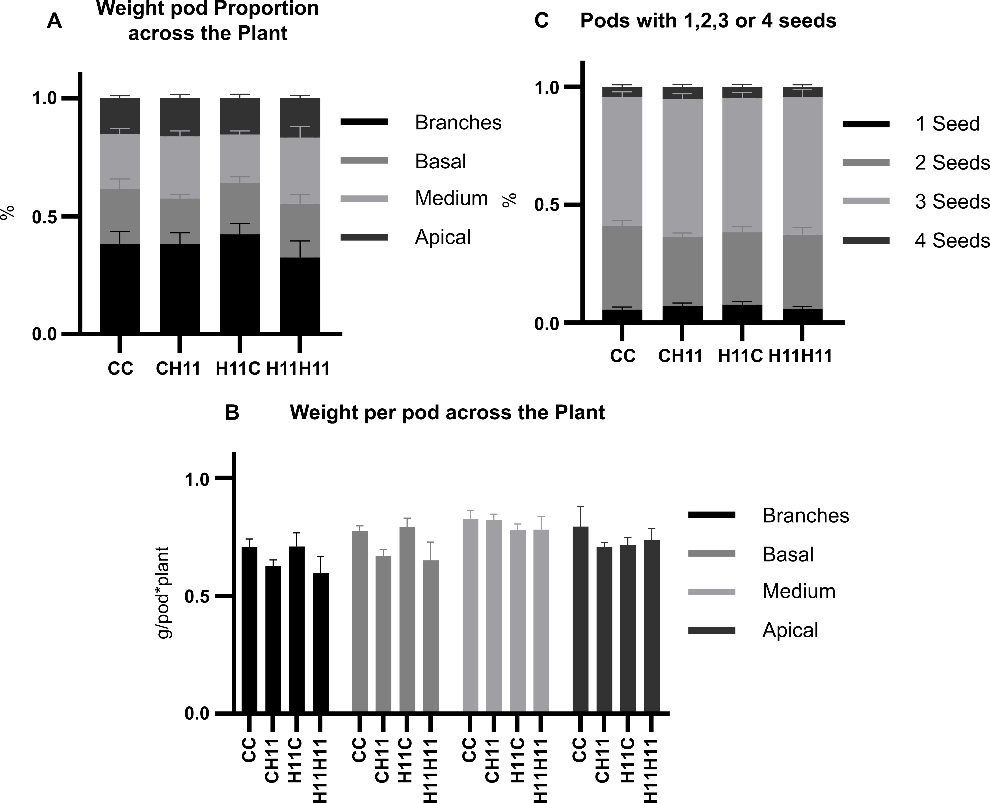

**Supplementary Figure S3. H11 scion plants have lighter pods in the basal portion of the stem**

Distribution of pod biomass and individual pod weight in the plant. The stem was divided into basal, medial, and apical sections. Branches and stems were analyzed separately. The basal section is the most proximal to the roots. (A) Total pod biomass partition through the stem. (B) Proportion of pods with 1, 2, 3 or 4 grains/pod. (C) Individual pod biomass weight in branches and basal, medial and apical portions of the stem.

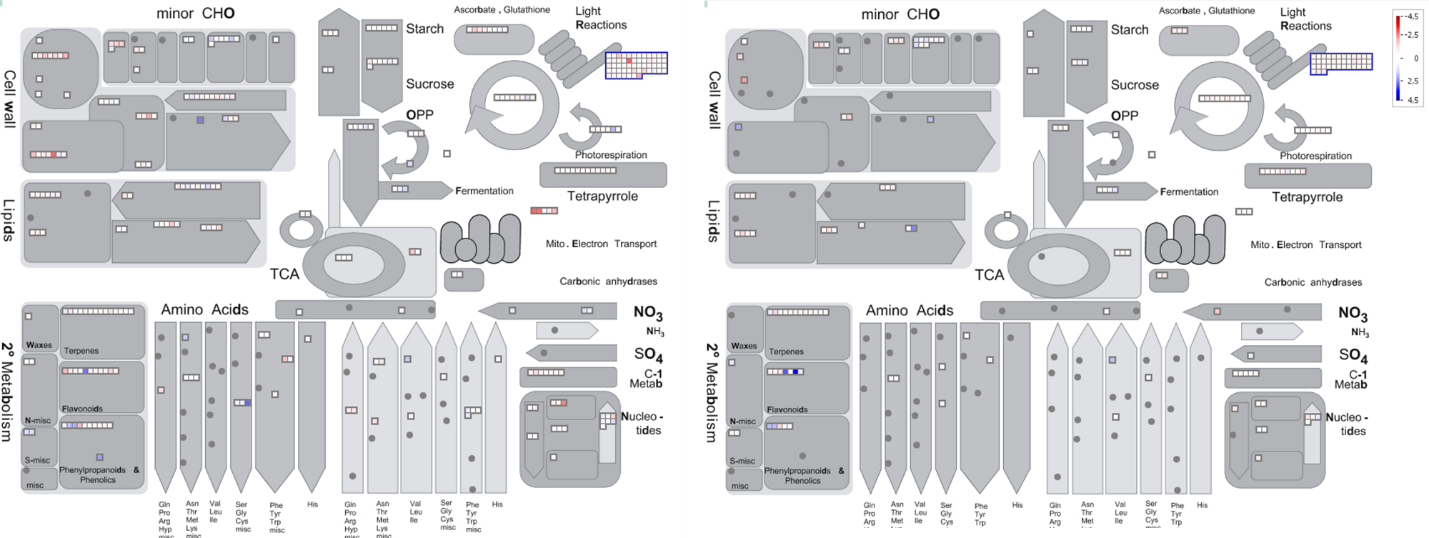

**Supplementary Figure S4:** **In the metabolism overview, H11 scion plants show repression of light-reaction genes when compared to CC plants**

MapMan metabolism overview of differentially expressed genes (DEGs). Left: H11H11 versus CC (homograft comparison); right: CH11 versus CC (heterograft versus homograft). Blue indicates repression and red, overexpression.

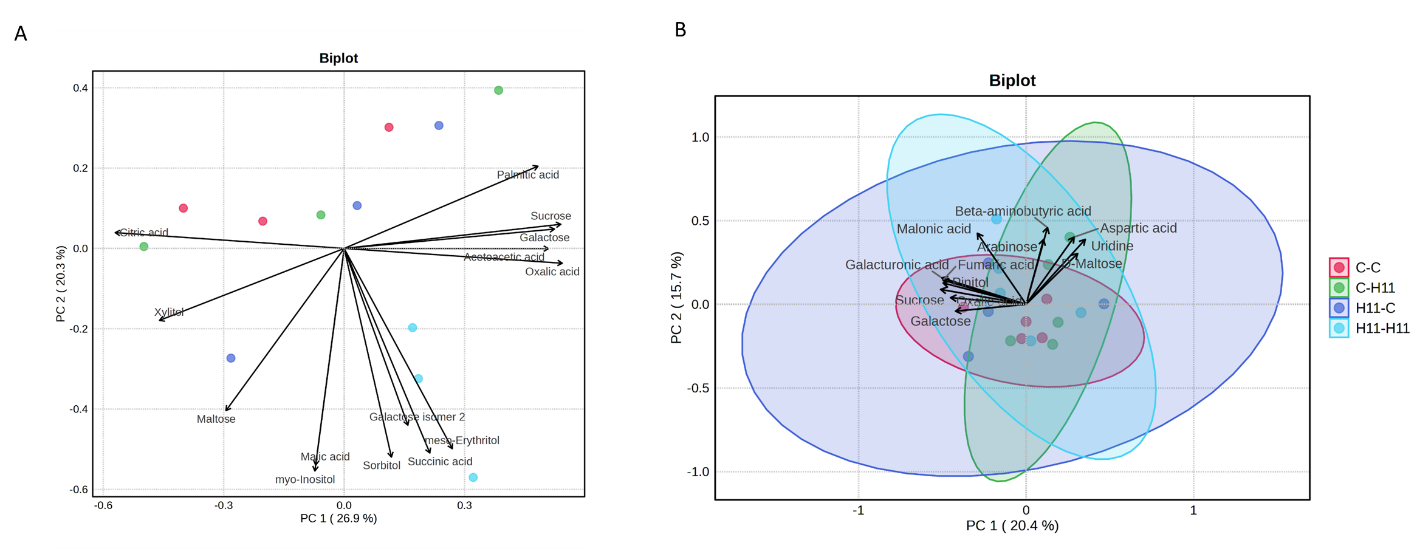

**Supplementary Figure S5. Principal component analysis of leaf metabolite profiles across grafting combinations.**

Biplots showing the first two principal components (PC1, PC2) of polar metabolite data measured by GC-MS in leaves of the four focal grafting combinations (CC, CH11, H11C, H11H11) at **(A)** solar noon (ZT8; n = 3 biological replicates per combination) and **(B)** end of day (ZT16; n = 4–5 per combination). Data were log₁₀-transformed and autoscaled before the PCA. Each dot corresponds to one biological replicate (a pool of leaves from three plants). Arrows indicate the loadings of the most informative metabolites. The percentage of variance explained by each component is indicated on the axes. 95 % confidence regions (shaded ellipses) are shown in (B) but not in (A), where the limited number of replicates per combination (n = 3) prevented their reliable estimation.

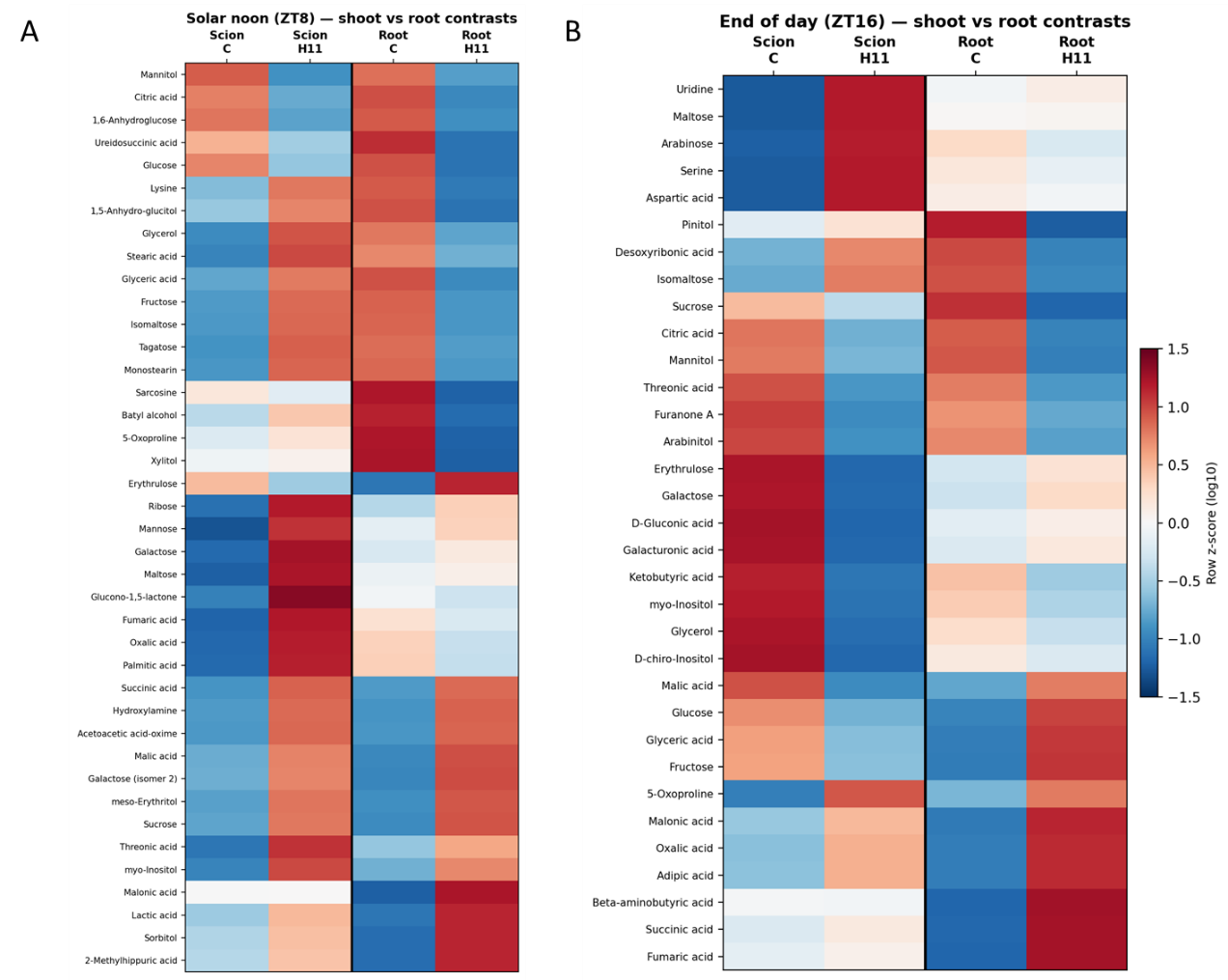

**Supplementary Figure S6. Leaf metabolite responses to scion and rootstock genotypes across grafting combinations**

Heatmaps showing the relative abundance of polar metabolites in leaves of the four focal grafting combinations, grouped according to scion or rootstock genotype, at **(A)** solar noon (ZT8) and **(B)** end of day (ZT16). For each metabolite, log₁₀-transformed values were averaged across all plants sharing a given genotype at the scion (Scion C: CC + H11C; Scion H11: CH11 + H11H11) or at the rootstock (Root C: CC + CH11; Root H11: H11C + H11H11) position; values are shown as row-wise z-scores of these group means, so colors are comparable across the four contrasts within a metabolite (row) but not between metabolites. The vertical black line separates the scion contrast (left) from the rootstock contrast (right). Rows are ordered by hierarchical clustering (Euclidean distance, average linkage).

**Supplementary Table S1. Comparison of homografted versus heterografted plants for yield-related traits**

Estimated marginal means (±SE) from linear models including graft type (homograft vs heterograft) and environment as fixed factors. Homografts: CC, H11H11, H4H4 (same genotype in scion and rootstock). Heterografts: all combinations with different scion and rootstock genotypes. P-values correspond to the heterograft-vs-homograft contrast. *** P<0.001, ** P<0.01, * P<0.05, P<0.10, ns P≥0.10. SE2 event excluded.

**Supplementary Table S2: Estimated marginal means (±SE) of yield-related traits per rootstock and scion genotype**

Estimated marginal means (±SE) of yield-related traits per rootstock or scion genotype, computed from linear models including Root, Shoot, and Environment (Env.) as fixed factors. Pairwise contrasts were performed against the control (C) using Dunnett's adjustment. *** P < 0.001, ** P < 0.01, * P < 0.05, P < 0.10, ns P ≥ 0.10.

**Supplementary Table S3. Effect of scion and rootstock genotypes on the progression of reproductive development in grafted soybean**

Plants were grouped by scion or rootstock genotype (root-scion notation). At each sampling date (40–121 days after sowing), the proportion of plants reaching or surpassing the indicated reproductive stage was compared among genotypes by χ² contingency test, with Benjamini-Hochberg FDR correction applied separately within each factor across sampling dates. For rows with a significant global test (q<0.05), different letters indicate genotypes that differ in pairwise Fisher's exact tests (FDR-corrected); genotypes sharing a letter do not differ. Where the global test was significant but pairwise comparisons did not resolve individual differences (Scion, 79 DAS), no letters are shown. Green: q<0.05.

**Supplementary Table S4: Vegetative and physiological traits at V3–V4 developmental stage**

Values are mean ± SE. No significant differences were detected among graft combinations for any of the evaluated traits (one-way ANOVA, p > 0.05). n = 5–7 plants per combination.

**Supplementary Table S5. Differentially expressed genes (DEGs) underlying the comparisons shown in Figure 5.**

Each column lists the DEGs of one set of the Figure 5 comparison. Genotypes follow the root-scion notation. Pairwise comparisons: CC-HH = CC vs H11H11; CC-CH = CC vs CH11; HH-HC = H11H11 vs H11C; HC-CH = H11C vs CH11. A vertical bar (|) denotes DEGs shared among the indicated comparisons; each gene appears once, under the exact set of comparisons in which it was differentially expressed. The comparison CC vs H11C yielded no DEGs and is therefore not represented. Gene identifiers follow the Glycine max genome Wm82.a4.v1.

**Supplementary Table S6: Significantly overrepresented Gene Ontology (GO) terms in pairwise RNA-Seq comparisons**.

GO enrichment analysis was performed in SoyBase using the differentially expressed genes (DEGs) from each pairwise comparison. Only Overrepresented terms with corrected P ≤ 0.05 are shown. BP: biological process; MF: molecular function; CC: cellular component.

**Supplementary Table S7: Pairwise Tukey HSD comparisons between graft combinations per metabolite at solar noon (ZT8)**

Group means ± SE are shown in a linear scale (ribitol-normalized peak area per mg fresh weight). For each pair of grafting combinations, the mean difference (group 1 − group 2) and the Tukey HSD-adjusted P-value are shown. Tests were performed on log₁₀-transformed data; P-values are corrected for multiple pairwise comparisons within each metabolite by the Tukey HSD procedure. Green shading: P < 0.05; yellow shading: P < 0.10. Metabolites are sorted by their lowest pairwise P-value, with significant ones (P < 0.05) appearing first.

**Supplementary Table S8: Pairwise Tukey HSD comparisons between graft combinations per metabolite at end of day (ZT16)**

Group means ± SE are shown in a linear scale (ribitol-normalized peak area per mg fresh weight). For each pair of grafting combinations, the mean difference (group 1 − group 2) and the Tukey HSD-adjusted P-value are shown. Tests were performed on log₁₀-transformed data; P-values are corrected for multiple pairwise comparisons within each metabolite by the Tukey HSD procedure. Green shading: P < 0.05; yellow shading: P < 0.10. Metabolites are sorted by their lowest pairwise P-value, with significant ones (P < 0.05) appearing first.

**Supplementary Table S9. Effect of scion and rootstock genotype on leaf polar metabolite abundance at solar noon (ZT8)**

For each metabolite, plants were grouped according to scion (C: CC + H11C; H11: CH11 + H11H11) or rootstock (C: CC + CH11; H11: H11C + H11H11) genotype, and the two groups were compared by Welch's t-tests on log₁₀-transformed data. Means ± SE in linear scale (ribitol-normalized peak area per mg fresh weight), fold change (FC = H11/C), p-value and Benjamini–Hochberg FDR (q) are shown for each contrast. Significance was considered at q < 0.05 (purple), with P < 0.05 (green) and P < 0.10 (yellow) reported as trends. Metabolites are sorted by their lowest p-value (scion or rootstock contrast); significant ones (P < 0.05) appear at the top.

**Supplementary Table S10. Effect of scion and rootstock genotype on leaf polar metabolite abundance at end of day (ZT16)**

For each metabolite, plants were grouped according to scion (C: CC + H11C; H11: CH11 + H11H11) or rootstock (C: CC + CH11; H11: H11C + H11H11) genotype, and the two groups were compared by Welch's t-tests on log₁₀-transformed data. Means ± SE in linear scale (ribitol-normalized peak area per mg fresh weight), fold change (FC = H11/C), p-value and Benjamini–Hochberg FDR (q) are shown for each contrast. Significance was considered at q < 0.05 (purple), with P < 0.05 (green) and P < 0.10 (yellow) reported as trends. Metabolites are sorted by their lowest p-value (scion or rootstock contrast); significant ones (P < 0.05) appear at the top.

**Supplementary Table S11. Pathway-level enrichment (MSEA, KEGG) of leaf metabolites affected by scion or rootstock genotype at ZT8 and ZT16**

Over-representation analysis of metabolites affected by the scion or the rootstock (P < 0.10, grouped t-tests on log10 data) at each time of day, performed with MetaboAnalyst against KEGG metabolite sets. Human KEGG pathways were used, as no plant-specific pathway library is available; central carbon metabolism is conserved across kingdoms. Only the two most informative organic-acid pathways are shown (TCA cycle and glyoxylate/dicarboxylate metabolism); redundant pathways supported by the same organic acids (e.g. alanine/aspartate/glutamate metabolism) and sugar pathways driven by free hexoses are not listed. Hits = input metabolites mapped to the pathway; n = total metabolites in the KEGG set. Supporting metabolites are listed for pathways with FDR < 0.05. Green: FDR < 0.05; yellow: 0.05 ≤ FDR < 0.10. FDR = Benjamini-Hochberg.

**Supplementary Table S12. Early physiological traits compared by scion genotype, rootstock genotype, and graft type**

Stomatal conductance (gsw), leaf temperature (Tleaf) and PSII quantum efficiency (ΦPSII) were measured with a LI-600 at the V2–V3 stage on the four graft combinations. For each variable, plants were grouped by scion (C: CC + H11C; H11: CH11 + H11H11), by rootstock (C: CC + CH11; H11: H11C + H11H11), or as homo- (CC + H11H11) versus heterografted (CH11 + H11C), and the two groups in each contrast were compared by Welch's t-test on the measured values. Group means and P-values are shown; the direction of each significant difference is indicated in parentheses. ** P < 0.01, * P < 0.05, P < 0.10, ns P ≥ 0.10.
